# Continuous psychophysics shows millisecond-scale visual processing delays are faithfully preserved in movement dynamics

**DOI:** 10.1101/2020.08.05.238642

**Authors:** Johannes Burge, Lawrence K. Cormack

## Abstract

Image differences between the eyes can cause interocular discrepancies in the speed of visual processing. Millisecond-scale differences in visual processing speed can cause dramatic misperceptions of the depth and 3D direction of moving objects. Here, we develop a monocular and binocular continuous target-tracking psychophysics paradigm that can quantify such tiny differences in visual processing speed. Human observers continuously tracked a target undergoing Brownian motion with a range of luminance levels in each eye. Suitable analyses recover the time-course of the visuomotor response in each condition, the dependence of visual processing speed on luminance level, and the temporal evolution of processing differences between the eyes. Importantly, using a direct within-observer comparison, we show that continuous target-tracking and traditional forced-choice psychophysical methods provide estimates of interocular delays that agree on average to within a fraction of a millisecond. Thus, visual processing delays are preserved in the movement dynamics of the hand. Finally, we show analytically, and partially confirm experimentally, that differences between the temporal impulse response functions in the two eyes predicts how lateral target motion causes misperceptions of motion-in-depth and associated tracking responses. Because continuous target-tracking can accurately recover millisecond-scale differences in visual processing speed and has multiple advantages over traditional psychophysics, it should facilitate the study of temporal processing in the future.

## Introduction

The binocular visual system combines spatial and temporal information from the eyes to estimate the structure of the three-dimensional environment, the three-dimensional motion of objects in the environment, and self-motion through the environment. A large body of research has focused on how spatial differences in the left- and right-eye images (i.e. binocular disparities) drive the estimation of 3D structure and motion (Banks, Gepshtein, & Landy, 2004; Burge & Geisler, 2014; Cormack, Czuba, Knöll, & Huk, 2017; Cormack, Stevenson, & Schor, 1991; Cumming & DeAngelis, 2001; DeAngelis, Ohzawa, & Freeman, 1991; Iyer & Burge, 2018; Julesz, 1964; Ogle, 1952; Ohzawa, DeAngelis, & Freeman, 1990; Tyler & Julesz, 1978; Wheatstone, 1838). A smaller but still substantial body of research has investigated how temporal differences in the processing of left- and right-eye images impact the estimation of motion-in-depth (Burge, Rodriguez-Lopez, & Dorronsoro, 2019; Carney, Paradiso, & Freeman, 1989; Lages, Mamassian, & Graf, 2003; Lit, 1949; Morgan & Thompson, 1975; Pulfrich, 1922; Reynaud & Hess, 2017; Rodriguez-Lopez, Chin, & Burge, 2023; Rogers & Anstis, 1972; Wilson & Anstis, 1969). Despite this long-standing interest (Watson, 1986), there have been few psychophysical attempts to measure the time course of visual processing and how it impacts the perception of motion in depth.

Traditional psychophysical data collection techniques, which require an observer to view a series of individual trials and to respond to each trial with a binary choice, usually do not reveal continuous temporal information about visual processing. Although recovering such information is possible in principle, traditional techniques are slow. Collecting sufficient data to recover time-course information is therefore often impractical. More generally, the slow pace of data collection places strong constraints on the number of conditions that can be practically run on each subject, and hence on the scope of the experimental questions that can be practically addressed using within-subjects designs.

Continuous target-tracking psychophysics, a new stimulus-response data collection technique, simultaneously provides information about the time course of visual processing and improves the rate of data collection (Bonnen, Burge, Yates, Pillow, & Cormack, 2015). With this technique, a property of a stimulus (e.g., position) is tracked via an input device (e.g. a computer mouse).^1^ Changes in the stimulus property of interest (e.g. contrast) are then associated with commensurate changes in the tracking response. The manner in which performance changes with the stimulus allows one to make inferences about various aspects of spatial and—as we show here—temporal processing.

There are many advantages of target-tracking psychophysics. First, as we will show, target tracking and traditional forced-choice psychophysics make near-identical estimates of temporal processing delays. Second, target tracking provides time-course information that traditional forced-choice psychophysics typically does not; this additional information provides a richer picture of the temporal characteristics of the system and can lead to enhanced predictive power (see Discussion). Third, target tracking provides a direct continuous measure of behavioral performance; other non-invasive continuous measures such as EEG and ERG (see for example, Kremers et al., 2022) provide more direct measures of neural activity, but must be related to performance via other means. Fourth, target tracking is low-tech and does not need specialized equipment other than a display screen and a mouse. (If target tracking in depth is desired, then a stereo-capable display is required). Fifth, target tracking requires little instruction or practice, and is therefore suitable for collecting large amounts of high-quality data from virtually any observer; this last advantage may make target tracking particularly advantageous for work with developmental, clinical, and other non-traditional populations.

The main goal of this paper is to demonstrate the utility of continuous psychophysics for measuring the time course of visual processing. To do so, we estimate interocular differences in visual processing using three methods—monocular target-tracking psychophysics (Fig. 1A), traditional forced-choice psychophysics (Fig. 1B), and binocular target tracking in depth (Fig. 1C)—and then compare results across methods.

**Figure 1.**
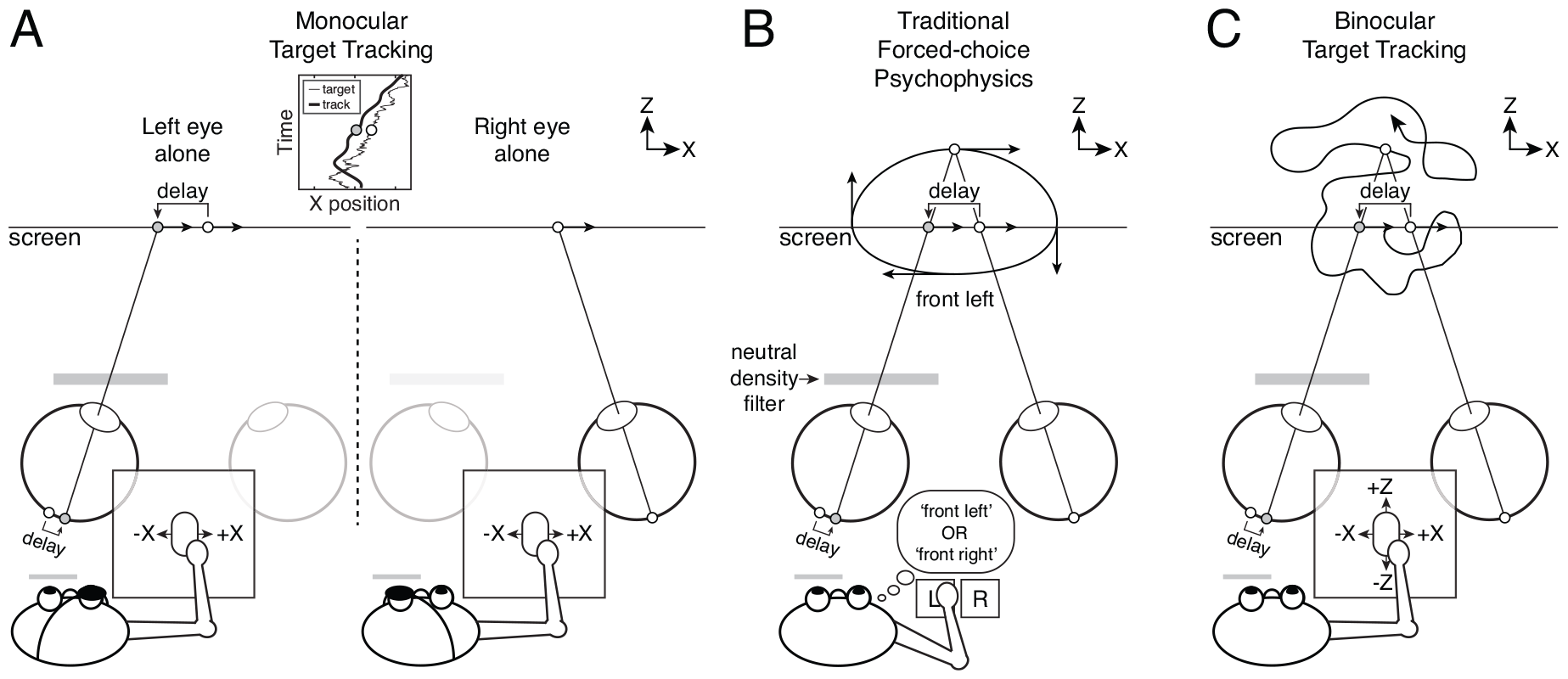
The classic Pulfrich effect, and three approaches to measuring interocular delays. **A** Monocular target tracking. A target undergoing a random walk in X on the screen is tracked while viewed with the left eye alone and also while viewed with the right eye alone. Although the target is always perceived in the plane of the screen (i.e. no illusory depth is perceived), comparing monocular tracking performance between the eyes can yield estimates of interocular delay that match those obtained with traditional psychophysics. **B** Traditional forced-choice psychophysics. When a moving target is viewed with an unequal amount of light in the two eyes, the distance and 3D direction of horizontal motion is perceived incorrectly. With a neutral density filter in front of the left eye, sinusoidal target motion (i.e. motion similar to that of a pendulum) in the screen plane is misperceived as motion along a near-elliptical trajectory in depth. The image in the darker eye is processed with a delay relative to the brighter eye. For rightward motion, this interocular delay causes the effective target image position in the darker eye (gray dot) to lag behind (i.e. to be shifted leftward) relative to the target image position in the brighter eye (white dot). For leftward motion, the target image position in the darker eye is shifted rightward (not shown). The binocular visual system computes the disparity from these effective left- and right-eye images, and the target is perceived behind the screen for rightward motion (and in front of the screen for leftward motion). In a traditional forced-choice psychophysical experiment, observers report their percept (e.g. ‘front left’ or ‘front right’) with a button press. This method recovers estimates of interocular delay. **C** Binocular target tracking. A target undergoing a random walk in X and Z is tracked while being viewed with both eyes. When the left eye is dark, the interocular delay causes a target moving rightward to be perceived farther away than it is and vice versa. Analyzing how target motion in X effects response motion is Z can also reveal interocular differences in processing.

To develop a stringent test for monocular target-tracking, we make use of the Pulfrich effect, a well-known stereo-motion phenomenon (Lit, 1949; Pulfrich, 1922). When the image in one eye is darker than the image in the other, motion in the frontal plane, like that of a clock pendulum, is misperceived as near-elliptical motion in depth (Fig. 1B). The effect occurs because the image with less light is processed more slowly. The interocular mismatch (i.e. delay) in processing speed causes, for moving objects, an effective neural disparity which leads to illusory percepts of depth. Interocular delays as small as a few milliseconds can cause large perceptual effects. Traditional forced-choice experiments which would, in this case, require observers to report whether a target stimulus was moving leftward or rightward when it appeared to be in front of the screen (see Fig. 1B) can accurately estimate these millisecond differences in processing speed. The Pulfrich effect therefore provides a demanding test for continuous psychophysics.

We show that continuous target-tracking psychophysics can be used to quantify millisecond-scale differences in visual processing between the eyes. We also prove a novel relationship between the temporal differences between left- and right-eye monocular processing and key properties of binocular target tracking in depth. These results indicate that sensory delays in visual processing are faithfully preserved in the movement dynamics of the hand. Target tracking has exquisite temporal sensitivity, provides estimates that are matched to those provided by traditional psychophysics with millisecond accuracy, and holds promise for a range of scientific and clinical applications.

## Results

### Monocular target-tracking psychophysics

First, we estimated the visual processing delays caused by luminance reductions with target tracking under monocular viewing conditions. Human observers tracked a white vertical target bar undergoing a random walk in X (i.e. horizontally) with a small mouse cursor dot (Fig. 2A). The task was to follow the target as accurately as possible. Tracking was performed monocularly under six viewing conditions: left eye alone or right eye alone at each of three different luminance levels (see Methods). The task was performed without difficulty; the human tracking response was a smoothed and delayed approximation of the target motion (Fig. 2B).

**Figure 2.**
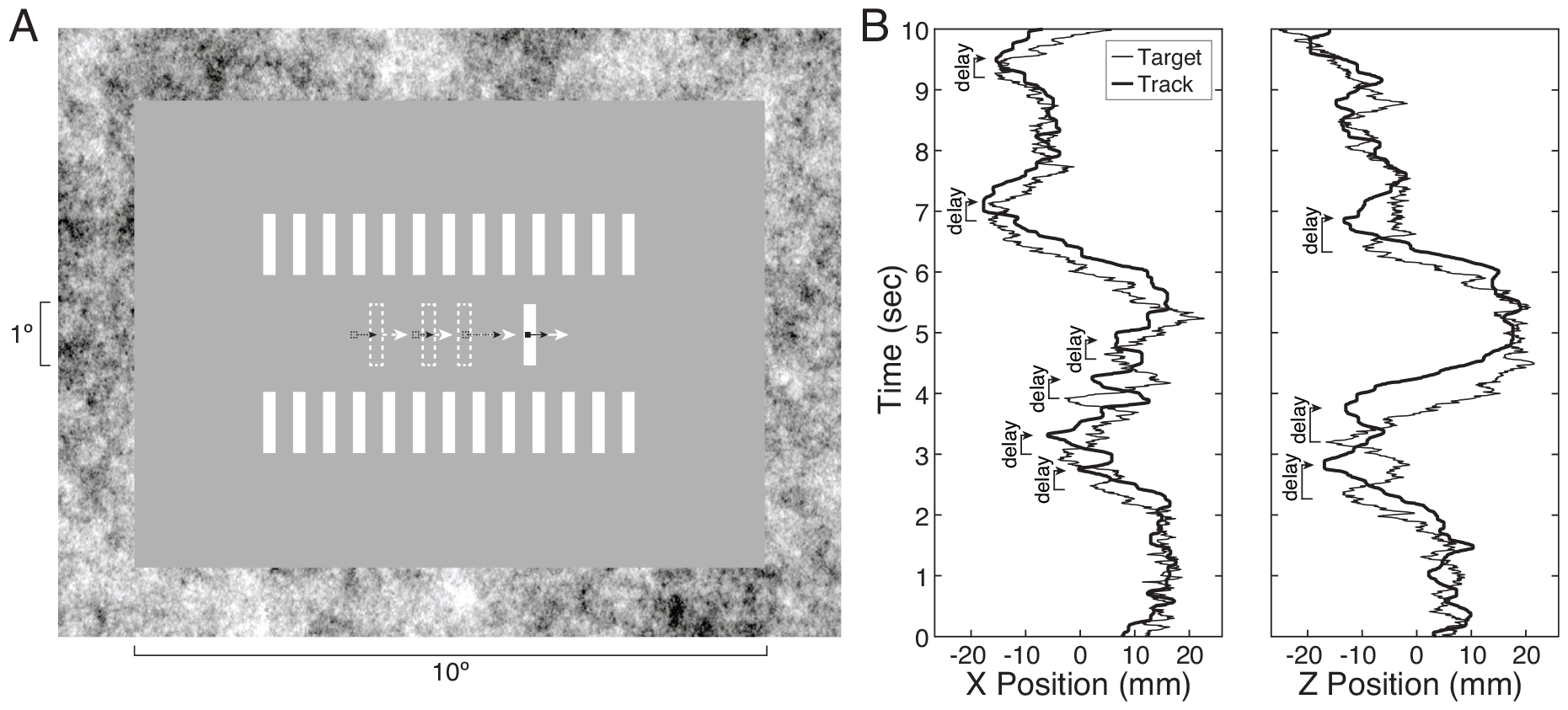
On-screen display and monocular tracking performance. **A** Target tracking stimulus. The target bar underwent a horizontal random walk in X or a random walk in both X and Z, depending on the experiment. The task was to track the target with a small dark mouse cursor. Motion direction and speed are indicated by arrows and dashed shapes; note that they are for illustrative purposes and were not present in the actual stimuli. **B** Target trajectory in X (solid curve) and tracked trajectory (dashed curve) across time. The response trajectory is a smoothed, delayed version of the target trajectory. Note that the delay in the human tracking response is approximately constant throughout the trial. **C** Target and response trajectories in Z. Note that the tracking delays are more pronounced in Z than in X.

Monocular cross-correlograms—the cross-correlation of the target and response velocities—are shown for all observers in the highest and lowest luminance conditions (Fig. 3A). The cross-correlogram approximates the shape of the temporal impulse response function of the visuomotor system, assuming the system is linear. The latency of the initial response, as quantified by the first point of the cross-correlogram that rises out of the noise, ranges between 150-200ms. The rise is steep such that the peak correlation occurs 50-75ms after the cross-correlation exceeds the correlation noise (see Methods). The temporal integration period ranges between 100-300ms across observers, as quantified by bandwidth (i.e. full-width at half-height). Note that because the motion statistics were matched across visual conditions, motor noise should be constant across conditions (Harris & Wolpert, 1998). (Jerk, the time derivative of response acceleration, does not vary systematically across conditions.).^2^ More systematic aspects of motor response should also remain constant across visual conditions. So, although the visuomotor response in any given condition will be affected by the characteristics of both the visual system and the motor system, comparisons across visual conditions should reveal changes in visual processing only.

**Figure 3.**
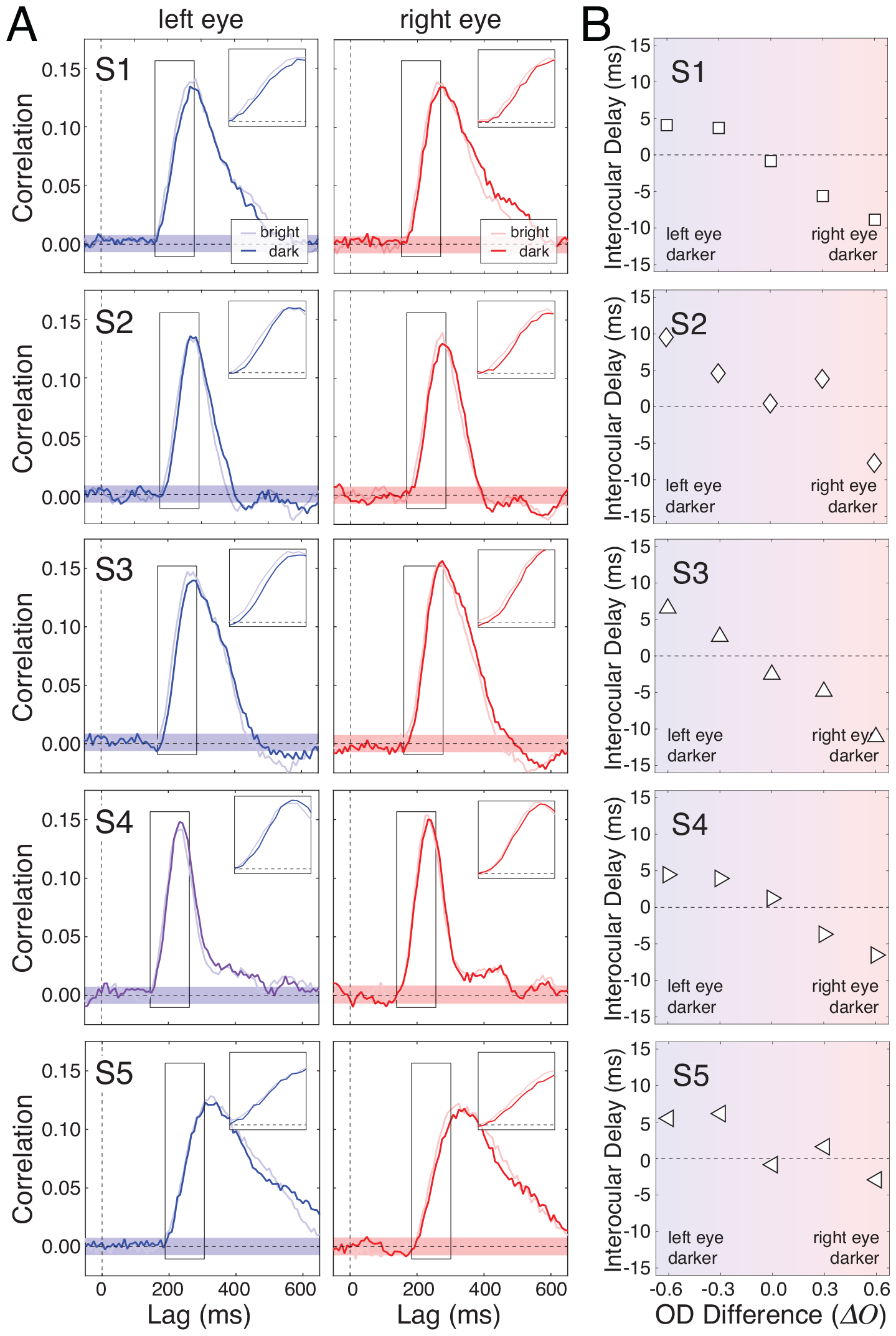
Monocular target tracking results. **A** Monocular target tracking in X for all observers. The cross-correlograms in the left column reflect left-eye-alone target tracking performance when the image was bright (i.e. maximum luminance: *OD*=0.0; light blue curve) and when the image was dark (i.e. one-quarter maximum luminance: *OD*=0.6; dark blue curve). The cross-correlograms in the right column similarly reflect right-eye-alone tracking performance when the image was bright and when the image was dark (light and dark red curves, respectively). The cross-correlograms are nearly overlapping. However, the dark curves are shifted rightward by a small but consistent amount. This shift indicates increased processing delays. Insets show the systematic delay between the rising edges of the cross-correlograms. The shaded bands in the main panel shows +1SD of the correlation noise, computed from lags less than 0ms. **B** Interocular tracking delays as a function of optical density difference. Negative optical density differences (*ΔO*=-0.6 and -0.3) show delays that were computed between conditions with dark left-eye images and bright right-eye images. Positive differences (*ΔO*=+0.3 and +0.6) show delays that were computed between conditions with bright left-eye images and dark right-eye images. In general, delay is systematically related to the interocular difference in optical density. (Please see Fig. 6A for error bars on these data.)

On quick examination, the monocular cross-correlograms appear to be the same for the high-and low-luminance conditions. Closer examination, however, reveals a small but systematic shift between the cross-correlograms in the two luminance conditions. This shift is clearest and most consistent in the rising edges of the cross-correlograms (Fig. 3A, insets). The visuomotor responses in the high luminance conditions are faster by several milliseconds than the responses in the low luminance conditions. Thus, assuming that the dynamics of the motor system itself are unchanged by the visual stimulus, the results imply that the speed of visual processing is faster in high than in low luminance conditions.

Figure 3B shows these differences in processing speed (i.e. delays) as a function of the luminance difference between the left and right eyes (i.e. simulated optical density difference; see Methods). We estimated the delays by computing the cross-correlation of the cross-correlograms in two luminance conditions (e.g. high luminance vs. low luminance); the delay yielding the maximum correlation was taken as the estimate of relative delay. This method makes use of all of the raw data, is robust, and is relatively unladen by assumptions. Across all observers, lower luminance caused slower visuomotor processing. In the conditions with the largest luminance differences, the delays ranged between 4 and 10 milliseconds. Interestingly, the estimated delays reported in Fig. 3B are in general agreement with previous psychophysical and neurophysiological investigations of the Pulfrich effect, which have shown that every log-unit reduction of luminance reduces the speed of processing by approximately 10 milliseconds (Carney et al., 1989; Rogers & Anstis, 1972). Moreover, the fact that these results are systematic and regular suggests that continuous target tracking is sensitive enough to estimate relative delays on the order of milliseconds.

The question, however, is whether these measured differences in visuomotor processing speed reflect differences in *visual* processing speed that are associated with stimuli of different luminance. To answer this question, we examined how estimates of temporal processing delays based on target tracking are related to estimates of temporal processing delays obtained with traditional forced-choice techniques. With forced-choice techniques, observer responses do not meaningfully depend on the dynamics of the motor system, so a quantitative comparison of delays in the two paradigms is a stringent test of whether target tracking can be used to measure millisecond-scale differences in visual processing.

### Traditional forced-choice psychophysics

To directly compare estimates of processing speed from tracking and traditional psychophysics, we used a standard paradigm to measure interocular delays associated with the Pulfrich effect (Burge et al., 2019; Rodriguez-Lopez, Dorronsoro, & Burge, 2020). The luminance levels in each eye were matched to those used in the monocular tracking experiment. The primary differences between the experiments were the motion trajectories followed by the stimulus and the method of response. Rather than a random walk, the target stimulus followed a sinusoidal trajectory on each eye’s monitor (Fig. 4ABC). Rather than continuously tracking the target stimulus with a cursor, observers viewed the target and made a binary response reporting an aspect of the 3D motion percept. When luminance differences between the eyes were non-zero, the target appeared to follow a near-elliptical 3D motion trajectory in depth.

**Figure 4.**
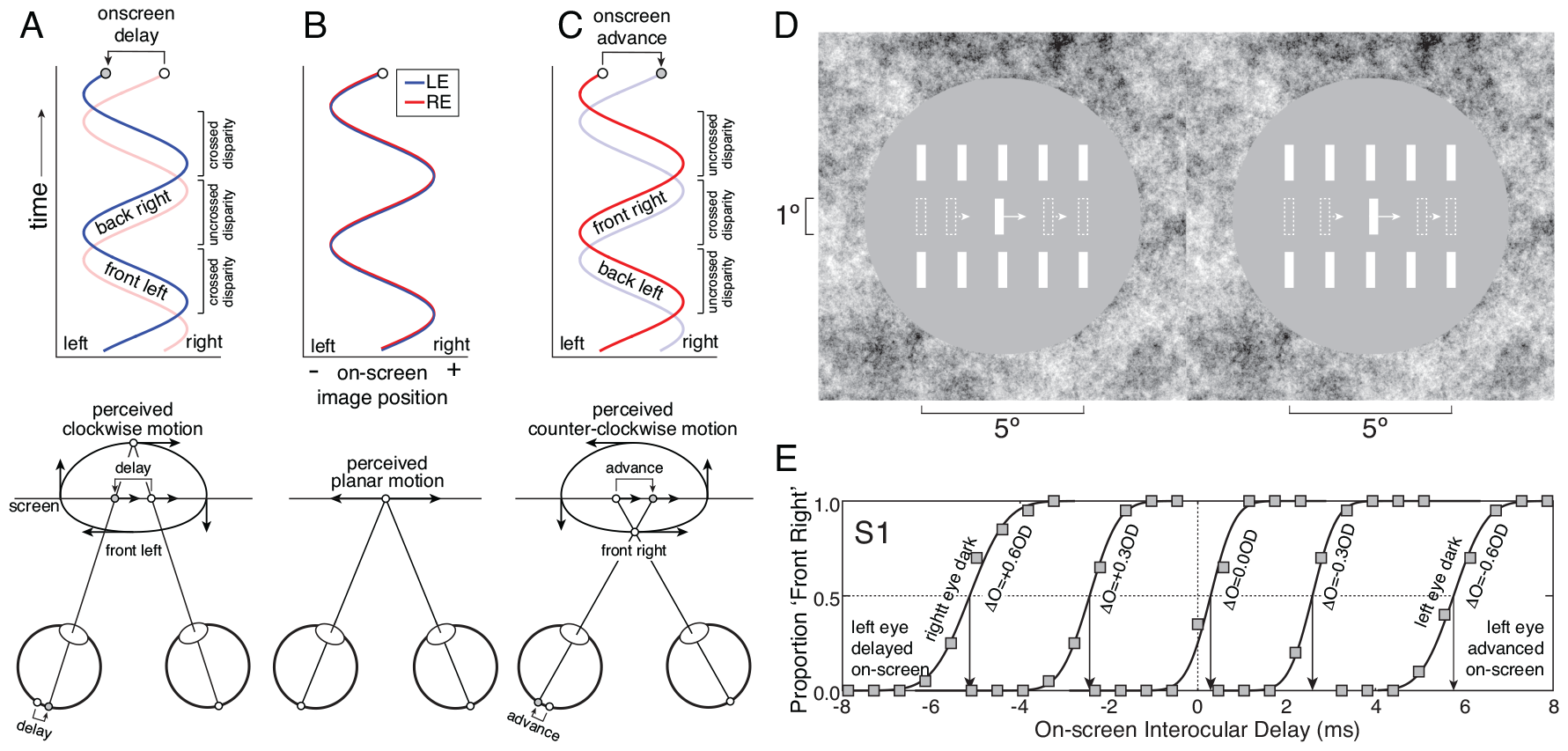
Binocular target stimulus, on-screen positions, and disparity-specified 3D target trajectories for forced-choice psychophysics experiment. **A** On-screen target image positions over time for the left and right eyes. When the left-eye image is delayed on-screen relative to the right-eye image (top), disparity specifies ‘front left’ 3D motion (bottom). **B** When no on-screen delays are present, disparity specifies motion in the plane of the screen. **C** When the left-eye image is advanced on-screen relative to the right-eye image, disparity specifies ‘front right’ 3D motion (bottom). **D** Binocular target stimulus. Arrows and dashed bars show motion direction and speed; they are for illustrative purposes and were not present in the actual stimuli. Free-fuse to see stimulus in 3D. Cross-fusers will see a depiction of ‘front right’ (i.e. counter-clockwise) 3D motion. Divergent-fusers will see a depiction of ‘front left’ (i.e. clockwise) 3D motion. **E** Example psychometric functions from the first human observer for interocular differences in optical density: *ΔO*=[-0.6OD, -0.3OD, 0.0OD, +0.3OD, +0.6OD]). These differences in optical density correspond to luminance differences ranging from the left eye having 75% less light than the right eye to the right eye having 75% less light than the left eye. To cancel the induced neural delays, the required on-screen interocular delays (arrows) change systematically with the luminance differences.

To change the stereoscopically-defined 3D motion trajectory, on-screen disparity was manipulated to simulate on-screen interocular delay (see Methods). To simulate the left-eye image being delayed on-screen relative to the right-eye image, on-screen disparity specified that the target was undergoing ‘front left’ motion (i.e. clockwise motion when viewed from above; Fig. 4A). To simulate no delay, disparity specified that the target was moving in the plane of the screen (Fig. 4B). And to simulate a left-eye image that is advanced on-screen relative to the right-eye image, on-screen disparity specified that the target was undergoing ‘front right’ motion (i.e. counter-clockwise motion when viewed from above; Fig. 4C). The size and shape of the target bar, the size and shape of the “picket fence” reference bars, the luminance levels, the peripheral 1/f noise were identical to those presented in the tracking experiment (Fig. 4D). The only differences were the shape of the mean luminance gray region (circular vs. rectangular) and the number of picket fence bars (five vs. thirteen).

The task was to report whether the target appeared to be moving leftward or rightward when it appeared to be in front of the screen. For a given interocular luminance difference, we measured a psychometric function: the proportion of trials that observers reported ‘front right’ as a function of the on-screen interocular delay (Fig. 4E). See Methods for details.

The goal of the experiment was to find, for each luminance difference, the critical on-screen delay that makes the target appear to move in the plane of the screen. This critical on-screen delay— the point of subjective equality (PSE) on the psychometric function—should be equal in magnitude and opposite in sign to the neural delay caused by the luminance difference between the eyes in each visual condition. We found that the critical on-screen delays (i.e. the PSEs) required to null the neural delays change systematically with the associated luminance differences in all five observers (Fig. 5).

**Figure 5.**
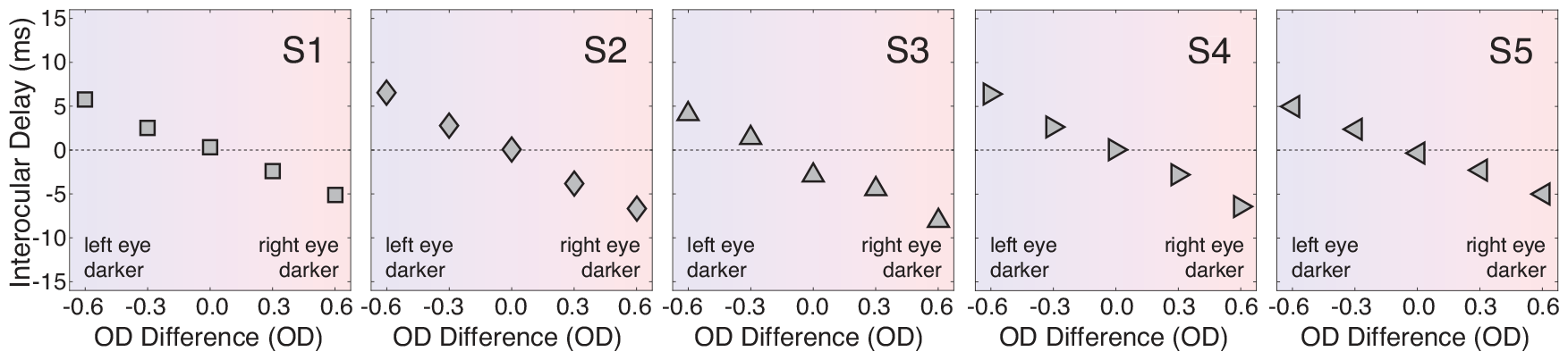
Traditional forced-choice psychophysics results. Critical on-screen interocular delays that resulted in percepts of zero motion in depth as a function of the interocular difference in optical density (i.e. *ΔO*=[-0.6OD, -0.3OD, 0.0OD, +0.3OD, +0.6OD]). These optical density differences correspond to the left-eye stimulus being 75% and 50% darker than the right-eye stimulus, the stimuli in both eyes having the same luminance, and the right-eye stimulus being 50% and 75% darker than the left-eye stimulus, respectively. Negative and positive on-screen delays respectively correspond to the left-eye stimulus being delayed and advanced relative to the right-eye on-screen stimulus. (Please see Fig. 6A for error bars on these data.)

### Target-tracking vs. forced-choice psychophysics

Figure 6A shows the estimates from the traditional forced-choice experiment plotted directly against the estimates from the continuous target-tracking experiment. Error bars in both directions are bootstrapped standard errors. The data are tightly clustered about the unity line, indicating millisecond-scale agreement between estimates of interocular delay provided by the two experiments (Fig. 6A). Across conditions and observers, the differences between the estimated delay with traditional and tracking psychophysics were very small. The mean difference in delay was -0.16ms (-1.04ms to 0.71ms; 95% confidence interval) with a standard deviation of 2.06ms (Fig. 6A, upper-right inset). The mean difference in delay was not statistically different from zero. For reference, Fig. 6B overlays the estimates of interocular delay from each of the two experiments (Note: Fig. 6B replots data from Figs. 3B & 5.)

**Figure 6.**
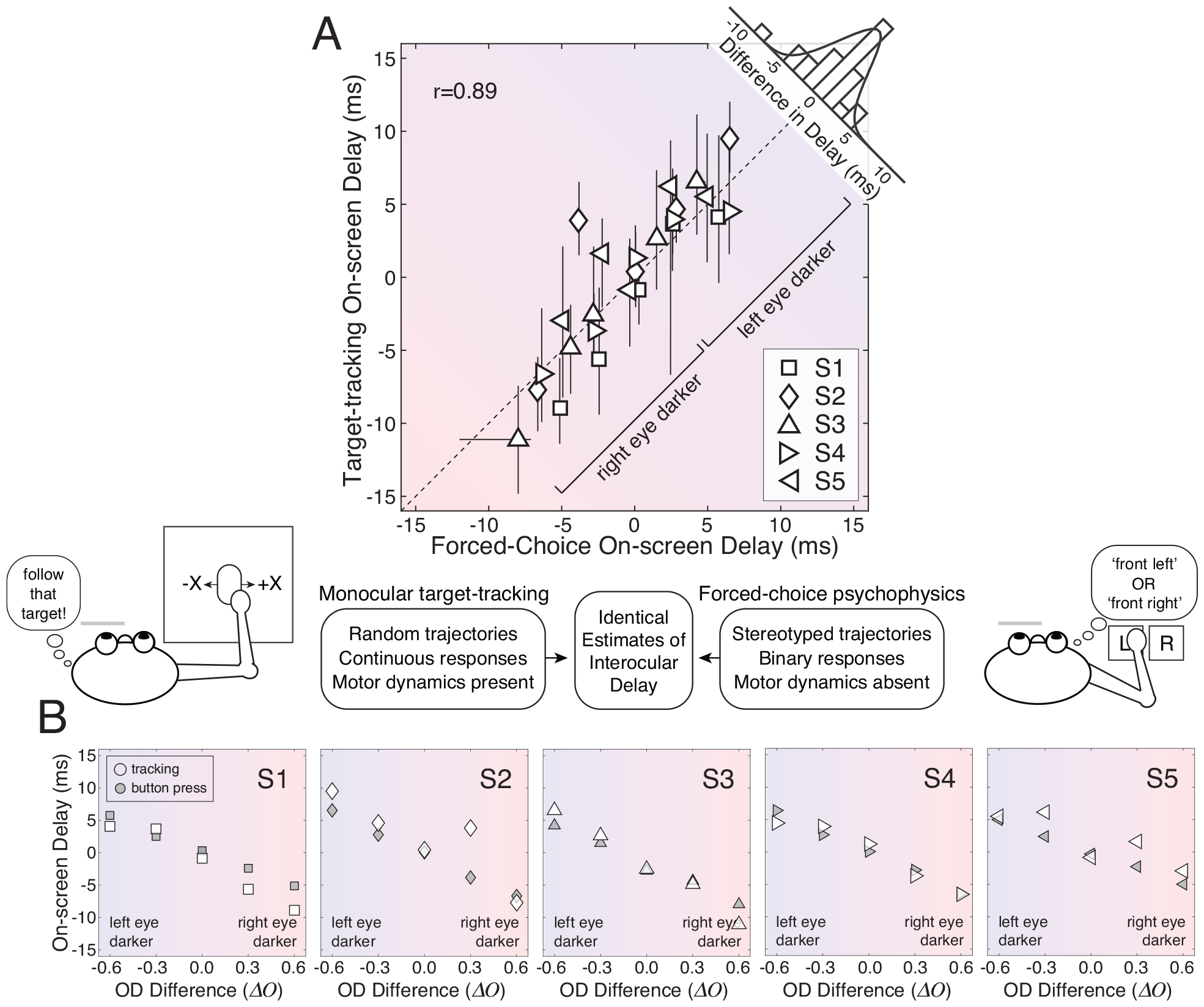
Interocular delays from target tracking vs. traditional forced-choice psychophysics. **A** Interocular delays measured with target tracking plotted against interocular delays measured with forced-choice psychophysics in equivalent conditions. The correlation is 0.89 (*p*=4.4x10^−9^). Red tint indicates conditions in which the right eye is darker. Blue tint indicates conditions in which the left eye is darker. Error bars indicate 68% bootstrapped confidence intervals from 1000 datasets resampled with replacement; if error bars are not visible, they are smaller than the size of the data symbol. The histogram (upper-right corner) shows the distribution of differences between the forced-choice delays and the tracking delays; the mean difference was only -0.16ms (SD=2.06ms). **B** Interocular delay as a function of interocular difference in optical density as measured with target tracking (white symbols) and forced-choice psychophysics (gray symbols) for individual observers in each of five different conditions (i.e. *ΔO*=[-0.6,-0.3,0.0,+0.3,+0.6]). Positive on-screen delays indicate that the left eye was processed more slowly. (Data replotted and overlayed from Figs. 3B & 5).

Given the enormous differences between the two psychophysical methods, the agreement between the respective estimates of interocular delay is striking. One method—traditional forced-choice psychophysics—presented a target stimulus following a stereotyped motion trajectory in depth (i.e. a near-elliptical path through a 3D volume of space) and obtained a binary response. This binary response reflected an aspect of the observer’s percept, and the response is completely independent of the temporal properties of the motor system. The other method— target-tracking psychophysics—presented a target stimulus following an unpredictable motion trajectory (i.e. a random walk in the 2D plane of the display monitor) and obtained the continuous motor response of the observer. This continuous response necessarily reflects the temporal properties of both the visual and motor systems. And yet, the estimates of interocular delay from the two methods agree to within a fraction of a millisecond on average, and to within a few milliseconds in each individual condition. The fact that substantially different stimuli and substantially different experimental paradigms yield near-identical estimates strongly suggests that both paradigms are measuring the same underlying quantity.

### Binocular target-tracking psychophysics

The preceding experiments show that monocular target tracking and traditional forced-choice psychophysics yield similar estimates of millisecond-scale processing differences between the eyes. Does binocular target tracking afford similarly precise measurements of interocular differences in visual processing? In this experiment, human observers binocularly viewed and tracked a target bar with a small cursor undergoing a random walk in the XZ plane (see Fig. 1C). Either the left- and right-eye on-screen images were equally bright or one eye’s image was substantially darker than the other (i.e. *ΔO*=[-0.6OD,0.0OD,0.6OD]). These luminance conditions were matched to those used in the monocular tracking experiment.

#### Binocular target-tracking: Empirical Results

Binocular tracking of horizontal target motion is similar to monocular tracking of horizontal motion for all luminance conditions (Fig. 7; X vs. X). The latency of the initial response ranges between 150-200ms across observers, and the temporal integration period ranges between 100-300ms. Binocular target tracking in depth is uniformly more sluggish (Fig. 7; Z vs. Z). In each observer, the latency of the initial response in Z occurs approximately 50ms later than the initial response in X, and the period of temporal integration in Z is nearly double the temporal integration period in X. These results hold when both eyes have the same luminance (Fig. 7A), when the left eye is darker than the right (Fig. 7B), and when the right eye is darker than the left (Fig. 7C). Hence, neither the X vs. X nor the Z vs. Z cross-correlograms provides information about differences in temporal processing between the eyes, as these cross-correlograms are near-identical down the columns of Fig. 7. The sluggishness of the response in Z to target motion in Z also replicates the primary finding from Bonnen et al. (2017) and is generally consistent with other results that have shown that changes in depth are processed more slowly than changes in horizontal position. The current results, however, do not shed light on the underlying reasons for the sluggishness of the Z motion processing.

**Figure 7.**
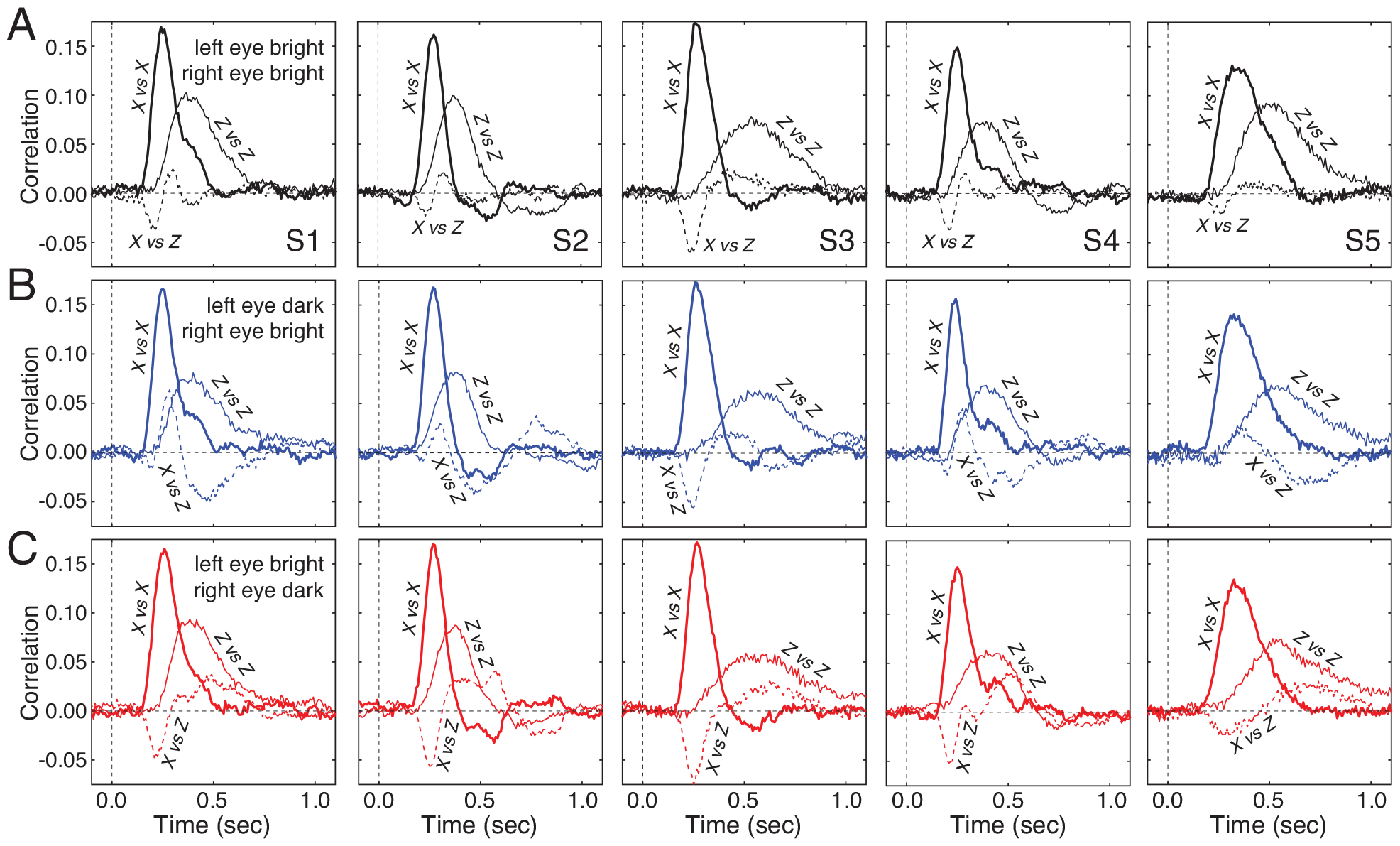
Binocular tracking. Temporal cross-correlograms between X target motion and X response motion (X vs. X), Z target motion and Z response motion (Z vs. Z), and X target motion and Z response motion (X vs. Z) for all five human observers for three visual conditions: **A** Both on-screen images are equally bright (*ΔO*= 0.0OD). **B** The left-eye image is dark (i.e. 75% of maximum luminance) and the right-eye image is bright (*ΔO*=-0.6OD). **C** The left-eye image is bright and the right-eye image is dark (i.e. 75% of maximum luminance; *ΔO*=+0.6OD). Tracking in X is comparatively swift, tracking in Z is more sluggish, and the impact of horizontal target motion on the depth response (X vs. Z) depends systematically on the luminance differences between the eyes.

Recall that, in the context of the Pulfrich effect (see Fig. 4), interocular processing delays cause horizontally-moving targets to be misperceived as moving in depth. We examined whether this signature of the Pulfrich effect is present in binocular tracking in depth; that is, we examined whether horizontal target motion is associated with response movements in depth. The cross-correlations between X target-motion and Z response-motion clearly show that such associations depend on the luminance condition (Fig. 7ABC; X vs. Z). When the left eye is dark (Fig. 7B), there tends to be an initial positive lobe, followed by a second negative lobe. When the right eye is dark (Fig. 7C), the shapes of the X vs. Z cross-correlograms are approximately mirror reversed. The dependence of the X vs. Z cross-correlograms on luminance condition suggests that they may be useful for recovering differences in the time-course of visual processing between the eyes. (Note that when both eyes are bright (Fig. 7A), there are small but systematic deviations from zero, which implies a baseline (*ΔO*=0.0OD) asymmetry in left- and right-eye processing; see below).

#### Binocular target-tracking: Model and Simulation

We developed a model to determine whether the X vs. Z cross-correlograms can be used to recover the time-course of differences in visual processing between the eyes. The model incorporates projective geometry, the left- and right-eye impulse response functions, and the statistics of target motion. We show analytically that the cross-correlation of X target-motion and Z response-motion is proportional to the difference between the temporal impulse response functions associated with the left and right eyes. That is

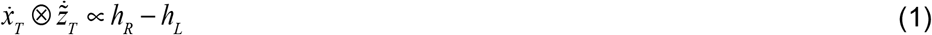

where 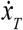 is the X-velocity of the target, 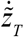 is the Z-velocity of the response, and *h*_*L*_ and *h*_*R*_ are the left- and right-eye impulse response functions, under the assumption of a linear system (see Supplement for derivation). Thus, it may be possible to use binocular 3D target tracking to estimate the temporal evolution of interocular differences in visual processing.

Simulations confirm the analytic result. The simulations were performed as follows. First, for a given virtual random walk in X and Z, we calculated the corresponding left- and right-eye on-screen image positions of the target on each time step using projective geometry. Next, we convolved the on-screen image positions with idealized left- and right-eye impulse response functions to obtain the left- and right-eye effective image positions. Lower luminance was assumed to shift the corresponding impulse response function in time. Finally, we back-projected the left- and right-eye effective image positions into 3D space (see Methods). The 3D tracking response was assumed to equal the position of the back-projected location in space.

Simulated tracking performance for an observer with different impulse response functions in the two eyes—one shifted to simulate the effect of lower luminance—is shown in Fig. 8. The impulse response functions are shown in Fig. 8A. The simulated observer smoothly tracks the horizontal target motion component with a consistent interocular delay, as in the monocular tracking experiment (Fig. 8B, top). Tracking performance in depth, however, is distinguished by obvious inaccuracies (Fig. 8B, bottom). The over- and under-shooting of the response in depth relative to the target depth depends systematically on horizontal target motion (Fig. 8B, rectangular boxes). This dependence is the hallmark of the Pulfrich effect, and can be quantified by the cross-correlogram of target X-motion with response Z-motion. The thick curves in Fig. 8C show this cross-correlogram. Importantly, the difference between the left- and right-eye temporal impulse response functions, shown by the thin curves in Fig. 8C, beautifully predicts the X vs. Z cross-correlogram, as indicated by Eq. 1.

**Figure 8.**
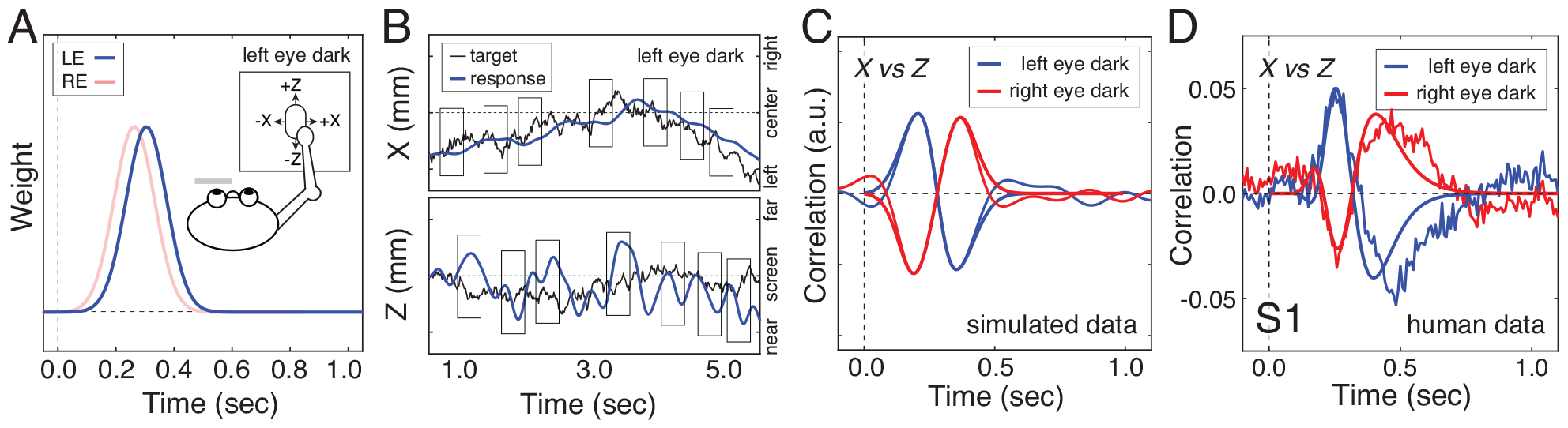
Simulated binocular tracking through depth with interocular luminance differences. **A** Impulse response functions for the model observer’s left and right eyes when the left eye is dark and the right eye is bright. **B** Resulting tracking performance in X and Z when the left eye is dark and processed with a delay relative to the right eye. Target motion in X is tracked smoothly and accurately, but large tracking inaccuracies are apparent in Z. Large over- and under-estimates of depth (i.e. Z position) are caused by rightward and leftward target movements, respectively, the hallmark of the classic Pulfrich effect. The vertical rectangles highlight regions of the time-series where these effects are most noticeable. **C** The cross-correlation of X target motion and Z response motion (thick curves) from the simulated data and the difference between the left- and right-eye impulse response functions (thin curves) are plotted for two different conditions with one eye dark and the other eye bright (colors). The simulated data agrees with the mathematical prediction in Eq. 1. **D**. Empirical (thin curves) vs. predicted (thick curves) data for the first human observer. The predictions are obtained by first fitting log-Gaussian functions to the cross-correlograms for relevant conditions in the monocular tracking experiment (see Fig. 3A), and then taking the difference of the fits (see Eq. 1).

#### Binocular target-tracking: Empirical results and model fits

To examine whether the empirical binocular target tracking data is predicted by discrepancies between the left- and right-eye impulse response functions, we fit the cross-correlograms in the bright and dark conditions of the monocular tracking experiment (see Fig. 3A) with log-Gaussian functions, took the difference of the fits (see Eq. 1), and compared the result to the empirically determined X vs. Z cross-correlograms. Data from the first human observer has clear similarities to the predictions (Fig. 8D, noisy vs. smooth curves).

Recall that the monocular left- and right-eye cross-correlograms reflect the processing dynamics of both the visual and the motor systems. However, if the dynamics of the motor system are stable across two monocular conditions (e.g. left eye dark, right eye bright), then the difference between the corresponding cross-correlograms should reflect only differences in visual processing (i.e. the differences between the left- and right-eye impulse response functions; see Eq. 1). Thus, assuming that non-linearities associated with binocular combination can be safely ignored in the context of this analysis (and no other complicating factors are present), the difference between the left- and right-eye cross-correlograms should predict the cross-correlogram between target X-motion and response Z-motion during binocular tracking, like it does in the simulated dataset.

The differences between fits to the monocular cross-correlograms (see Methods) nicely predict the binocular X vs. Z cross-correlograms in the first human observer (Fig. 8D, smooth curves). (The baseline X vs. Z asymmetry in the left-eye bright, right-eye bright condition was subtracted off before making the comparison; see Fig. 7A.) In this observer, binocular tracking performance is predicted by discrepancies between the left and right-eye impulse response functions. Thus, when the left- and right-eye on-screen images have different luminance levels, the impact of horizontal target motion on the human response in depth is well-accounted for by differences in the monocular cross-correlograms.

Across observers, however, the prediction quality is mixed, and in some cases it is poor (Fig. 9). The first lobes of the cross-correlograms are reasonably predicted in some observers but not at all in others. The fact that the initial lobes are well-captured in the first two observers is, however, intriguing. Additional lobes of the cross-correlograms are poorly accounted for in all observers (except perhaps for observer S1). (The prediction quality–and the raw data–is particularly bad for observer S3, who reported struggling to track the target in depth, although one would be hard-pressed to detect these struggles from the Z vs. Z cross-correlograms.) We speculate that these additional lobes, which tend to occur half a second or more after the onset stimulus motion are driven by fundamentally different processes (e.g. feedback delays due to binocular combination (Goldreich, Krauzlis, & Lisberger, 1992)) than those that are measured in the monocular tracking and traditional forced-choice psychophysics experiments. Hence, although XZ correlation data from the binocular conditions are well-suited, in theory, to estimate the time-course of temporal processing differences between the eyes (see Fig. 8A-C), the discrepancies between model and human cross-correlograms—especially after ∼500ms—make such analysis impractical with the present framework.

**Figure 9.**
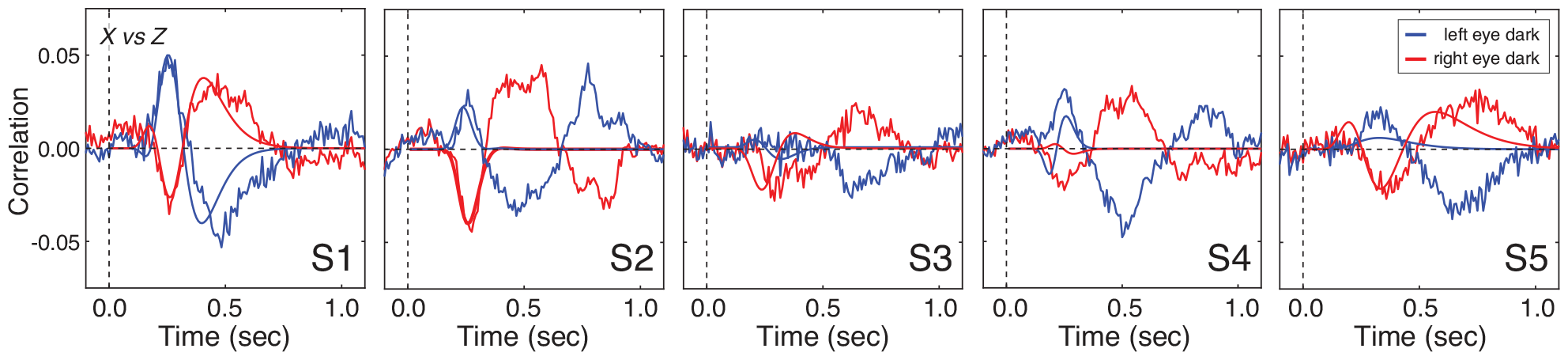
Binocular tracking in depth for all human observers (Note: data and predictions in the left-most panel are identical to the data and predictions in Fig. 8D). Horizontal target motion systematically affects human tracking responses in depth curves (thin, noisy curves). The blue and red curves depict how the X vs. Z cross-correlograms in the unequal luminance conditions (*ΔO*=-0.6 and *ΔO*=+0.6] the change from baseline (i.e. both eyes bright)(see Methods). Blue curves represent the X vs. Z cross-correlograms when the left eye is dark and the right eye is bright. Red curves represent the X vs. Z cross-correlograms when the left eye is bright and the right eye is dark. Predictions from the difference between the monocular cross-correlograms (thick, smooth curves) in the corresponding visual conditions (dark eye: *OD*=0.6, bright eye: *OD*=0.0) account for the first lobe of the binocular X vs Z cross-correlograms in roughly half of the conditions. The second lobes and third lobes of the cross-correlograms are poorly accounted for. The uneven ability of the model (see Eq. 1) to predict the data suggests that other unmodeled factors contribute to binocular tracking performance.

The analytical results, simulations, and partial experimental validation regarding tracking in depth are intriguing, but the discrepancies between the predictions and results suggest that more work must be done. The fact that the additional lobes are unaccounted for by the predictions suggests the involvement of other processes in binocular combination or in binocular tracking in depth. These phenomena are potentially quite interesting, but a full analysis of them must await targeted experiments and analyses in the future.

## Methods

### Participants

Five human observers participated in the experiment; four males and one female. Two were authors and the rest were naïve to the purposes of the experiment. The observers were aged 26, 31, 36, 41, and 55 years old at the time of the measurements. All had normal or corrected-to-normal visual acuity (20/20), and normal stereoacuity (20arcsec) as determined by the Titmus Stereo Test. And all provided informed consent in accordance with the Declaration of Helsinki using a protocol approved by the Institutional Review Board at the University of Pennsylvania.

### Apparatus

Stimuli were displayed on a custom-built four-mirror stereoscope. Left- and right-eye on-screen images were presented on two identical Vpixx VIEWPixx monitors (https://vpixx.com/products/viewpixx/). The gamma function of each monitor was linearized using custom software routines. The monitors were 52.2x29.1cm, with a spatial resolution of 1920x1080 pixels, a refresh rate of 120Hz, and a maximum luminance of 105.9cd/m^2^. After light loss due to mirror reflections, the maximum luminance was 93.9cd/m^2^. A single AMD FirePro D500 graphics card with 3GB GDDR5 VRAM controlled both monitors to ensure that the left and right eye images were presented simultaneously. To overcome bandwidth limitations of the monitor cables, custom firmware was written so that a single color channel drove each monitor; the red channel drove the left monitor and the green channel drove the right monitor. The single-channel drive to each monitor was then split to all three channels for gray scale presentation.

Observers viewed the monitors through a pair of mirror cubes positioned one inter-ocular distance apart (Supplementary Fig. S1). Heads were stabilized with a chin and forehead rest. The mirrors were adjusted such that the vergence distance matched the distance of the monitors. The monitors were positioned at a distance of 100cm. This distance was confirmed both by a laser ruler measurement and by a visual comparison with a real target at 100cm. The mirror cubes had 2.5cm openings which yielded a field of view of 15x15º. At this distance, each pixel subtended 1.09arcmin. Stimulus presentation was controlled via the Psychophysics Toolbox-3 (Brainard, 1997). Anti-aliasing enabled accurate presentations of disparities as small as 15-20arcsec.

### Neutral density filters

To control the stimulus luminance for each eye and to induce luminance differences between the eyes we placed ‘virtual’ neutral density filters in front of the eyes. First, we converted optical density to transmittance, the proportion of incident light that is passed through the filter, using the standard expression *T* = 10^−*OD*^ where *T* is transmittance and *OD* is optical density. Then, we reduced the monitor luminance by a scale factor equal to the transmittance. We have previously verified that real and virtual neutral density filters with equivalent strengths yield identical performance (Burge et al., 2019). The optical density of the neutral density filters took on values of 0.0, 0.3, and 0.6. For the experiments with binocular viewing conditions (binocular tracking and forced-choice psychophysics), I interocular difference in optical density

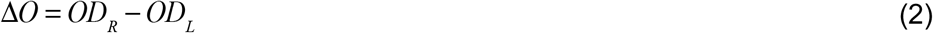

quantifies the luminance difference between the eyes. In all binocular conditions, at least one eye had an optical density of 0.0. Interocular differences in optical density thus ranged from -0.6 and 0.6 (i.e. *ΔO* = [-0.6, -0.3, 0.0, +0.3, +0.6]). At the extremes of this range, the stimulus to one eye had only 25% the luminance of the stimulus to the other eye.

### Procedure: Tracking experiments

Tracking data and forced-choice data was collected from each human observer in a within subjects design. In the tracking experiments, the observers controlled a mouse with their preferred hand in a comfortable position; all were right-handed. Observers initiated each run by clicking the mouse, which caused the target bar to appear in the center of the screen. After a stationary period of 500ms, the target followed a random-walk trajectory for the next eleven seconds. The task was to track the target as accurately as possible with a small dark mouse cursor.

In the monocular tracking experiment, data was collected in left-eye-alone and right-eye-alone conditions. For each eye, tracking data was collected at one-quarter maximum luminance, at half maximum luminance, and at maximum luminance. These values are equivalent to placing neutral density filters with optical densities of 0.6, 0.3, and 0.0, respectively, in front of the viewing eye. The non-viewing eye was occluded with an eye patch. Data was collected in twenty intermixed blocks of twelve runs each for a total 40 runs per condition.

In the binocular target tracking experiment, data was collected in each of three luminance conditions that were the analogs of the conditions in the monocular tracking experiment: left eye at one-quarter maximum luminance with the right eye at maximum luminance (i.e. left eye dark & right eye bright; *ΔO=*-0.6OD), both left and right eye at maximum luminance (*ΔO=*0.0OD), and left eye at maximum luminance and right eye at one-quarter maximum luminance (i.e. left eye bright and right eye dark; *ΔO=*+0.6OD). Data was collected in ten intermixed blocks of twelve runs each for a total of 40 runs per condition were collected.

Left-eye-alone monocular tracking blocks, right-eye-alone monocular tracking blocks, and binocular tracking blocks were interleaved and counterbalanced.

### Procedure: Forced-choice experiments

In the forced-choice experiment the observer’s task was to report, with a button press, whether the target bar was moving leftward or rightward when it appeared to be nearer than the screen on its virtual trajectory in depth. Using a one-interval two-alternative forced choice procedure, nine-level psychometric functions were measured in each condition using the method of constant stimuli. The levels of on-screen interocular delay ranged between +10.0ms at maximum. The range and particular levels were set according to the sensitivity with which each human observer discriminated interocular delay. This was done to ensure good sampling of the psychometric functions. The psychometric functions were fit with a cumulative Gaussian via maximum likelihood methods. The 50% point on the psychometric function—the point of subjective equality (PSE)— indicates the on-screen interocular delay needed to null the interocular difference in processing speed. Observers ran 180 trials per condition in counter-balanced blocks of 90 trials each.

### Stimuli: Tracking experiments

For the tracking experiments, the target bar was a moving white vertical bar on a gray background. The target bar, which subtended 0.25x1.00º of visual angle, was vertically flanked by two stationary sets of thirteen picket fence bars (Fig. 2A). The gray background subtended 10.0x7.5º of visual angle.

The target bar performed a random walk in X only or in X and Z. The X and Z positions of the target on each time step *t* + 1 were generated as follows

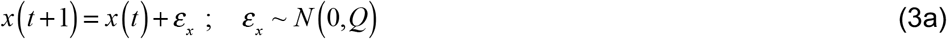

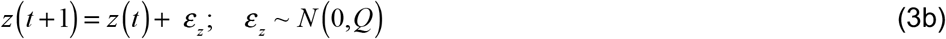

where *ε*_*x*_ and *ε*_*z*_ are samples of Gaussian noise that determine the change in target position from the current to the next time step, and *Q* is the so-called drift variance. The drift variance controls the mean magnitude of the position change on each time step, and hence the overall variability of the random walk. The variance of the walk positions across multiple walk instantiations *σ*^2^ (*t*) = *Qt* is equal to the product of the drift variance and the number of elapsed time steps, *t*. The value of the drift variance in our task (0.8mm per time step = 2.75arcmin per time step) was chosen to be as large as possible such that each walk would traverse as much ground as possible while maintaining the expectation that less than one walk out of 500 (i.e. less than one per human observer throughout the experiment) would escape the horizontal extent of the gray background area (176x131mm; 10.0x7.5º) before the 11 second trial completed. Ninety-five percent of the visited horizontal positions were within 50mm of the starting point in X and in Z. Ninety-five percent of the visited positions in depth corresponded to disparities smaller than 11arcm, and less than one walk out of 500 generated a disparity larger than 20arcmin. Even these largest disparities are much smaller than the upper disparity processing limit for stereopsis

In many experiments involving stereopsis, the distance of the virtual stimulus is specified by retinal disparity, which is directly determined by the stimulus position on each of two eye-specific monitors. The on-screen stimulus positions corresponding to a particular target position in stereoscopic space can be obtained using similar triangles (Supplementary Fig. S2). The left- and right-eye on-screen image positions corresponding to a virtual target position, are given by

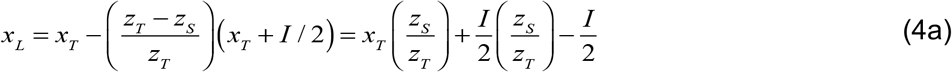

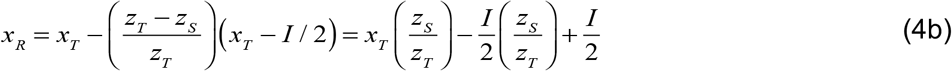

where *z*_*S*_ is the screen distance and *I* is the interocular distance.

### Model: Tracking response in 3D

The effective left- and right-eye images are obtained by convolving the on-screen left- and right-eye images with eye-specific temporal impulse response functions

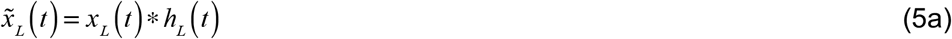

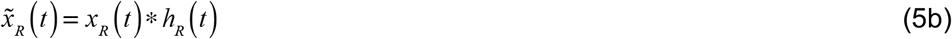

where *h*_*L*_ (*t*) and *h*_*R*_ (*t*) are the left- and right-eye temporal impulse response functions, respectively. Convolving the left- and right-eye target velocities with the impulse response function of each respective eye gives the velocities of the effective left- and right-eye images.

The predicted 3D response position is obtained by back-projecting the effective left- and right-eye images into 3D space. The X and Z positions of the 3D response are given by

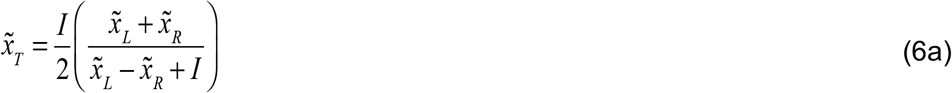

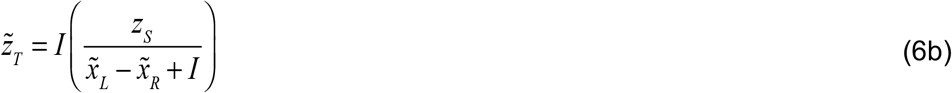

For the monocular tracking conditions, the stimuli only changed position in X and the observers wore an eye patch over one eye (see Fig. 1B).

The response velocities in X and Z are obtained by differentiating the X and Z response positions with respect to time

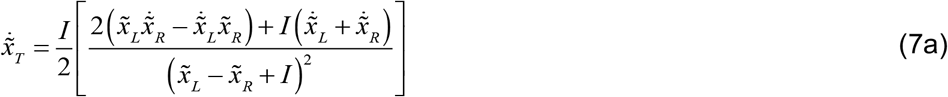

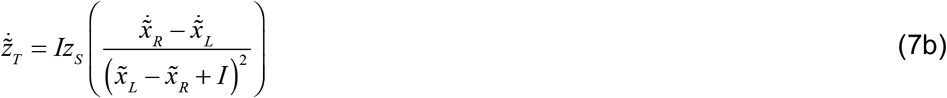

where 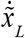 and 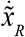 are the velocities of the effective target images in the left and right eyes, respectively. The target velocities in X and Z are given by functions having an identical form, provided that the velocities of the left- and right-eye on-screen images are substituted for the velocities of the effective target images in the two eyes.

To determine the impulse response function relating the target and response, we computed the zero-mean normalized cross-correlations between the target and response velocities

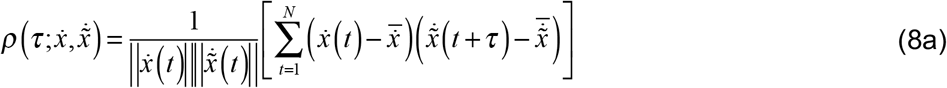

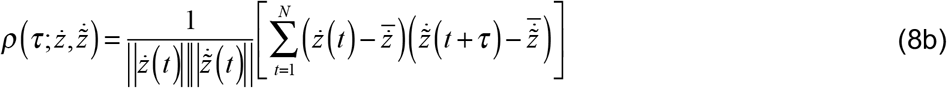

where *τ* is the lag, 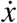 and 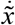 are the target and response velocities in X,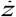 and 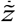 are the target and response velocities in Z. We also computed the zero-mean normalized cross-correlation between the X target velocity and the Z response velocity

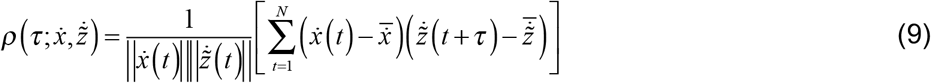

to determine how target X motion impacts response Z motion. The influence of X target motion on Z response motion is the hallmark of the Pulfrich effect.

The normalized cross-correlation is obtained by performing a series of normalized dot products between two time-series at each of many different lags. It quantifies the similarity of two time-series as a function of time lag. Assuming a linear system, when the input time series (i.e. the target velocities) is white noise, as it is here, the cross-correlation with the response gives the impulse response function of the system. When computing the normalized cross-correlations, we excluded the first second of each eleven second tracking run ensure only steady state performance is analyzed. (The time-series data indicate that most subjects had acquired the target within the first 500 milliseconds of each trial.) First, we computed the normalized cross-correlation in each run (Eqs. 8 & 9). Then, we averaged these cross-correlograms across runs in each condition. It is these mean cross-correlation functions (i.e. the cross-correlograms) that are presented in the figures.

### Predicting binocular from monocular tracking performance

Predicting binocular tracking performance from monocular tracking performance was a three-step process. To determine how lateral target motion influences the depth response we first fit the monocular cross-correlograms with log-Gaussian shaped functions using least squared regression. In nearly all cases, the log-Gaussian functions provided excellent fits to the monocular cross-correlograms once the curves exceeded two standard deviations of the correlation noise, which was computed from lags less than zero milliseconds. Second, we computed the difference between the fits in the bright condition (*OD*=0.0) in one eye and the dark condition (*OD*=0.6) in the other eye (e.g. left eye dark, right eye bright). Third, we subtracted the X vs. Z cross-correlogram in the condition where both eyes are bright from the cross-correlograms in the conditions of interest (i.e. one eye dark, one eye bright). This subtraction removes each observer’s baseline asymmetry, and isolates the changes caused by the interocular luminance differences.

### Stimuli: Forced-choice experiments

For the forced-choice psychophysics, we simulated the classic pendulum Pulfrich stimulus on the display. For each trial, the left- and right-eye on-screen target positions in degrees of visual angle were given by

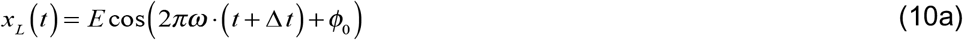

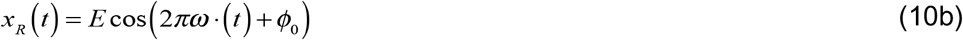

where *E* is the target movement amplitude in degrees of visual angle, *ω* is the temporal frequency of the target movement, *ϕ*_0_ is the starting phase, *t* is time in seconds, and Δ*t* is the on-screen delay between the left- and right-eye target images. When the interocular on-screen delay is non-zero, a spatial binocular disparity results, and the target follows a near-elliptical trajectory of motion in depth. Negative values indicate that the left-eye on-screen image is delayed relative to the right-eye image. Positive values indicate that the left-eye on-screen image is advanced.

Both eyes’ images were presented coincidently on each monitor refresh with an on-screen binocular disparity that was equivalent to the desired on-screen delay. We calculated the equivalent on-screen disparity 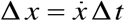 from the target velocity and the desired on-screen delay and then shifted the on-screen spatial positions of the left- and right-eye images. The equivalent on-screen disparity as a function of time is given by

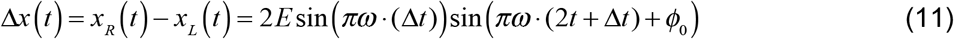

where negative disparities are crossed (i.e. nearer than the screen) and positive disparities are uncrossed (i.e. farther than the screen). The maximum disparity magnitude occurs when the perceived stimulus is directly in front of the observer and the lateral movement is at its maximum speed. When the stimulus is moving to the right, the maximum disparity in visual angle is given by Δ*x*_*max*_ = 2*E*sin(*π*ω).

The movement amplitude was 2.5º of visual angle (i.e. 5.0º total change in visual angle; see Fig. 4D), the temporal frequency was 1 cycle per second, and the starting phase *ϕ*_0_ was randomly chosen to be either 0 or *π*. Restricting the starting phase to these two values forced the stimuli to start either 2.5º to the right or left of center, respectively, on each trial.

Two sets of five vertical 0.25x1.00º bars in a ‘picket fence’ arrangement flanked the region of the screen traversed by the target bar (Fig. 4D). The picket fences were defined by disparity to be in the plane of the screen, and served as a stereoscopic reference of the screen distance. A 1/f noise texture, also defined by disparity to be in the plane of the screen, covered the periphery of the display to help anchor vergence and serve as a further stereoscopic reference to the screen distance. A small fixation dot was located at the center of the screen, at which observers were instructed to fixate.

## Discussion

We have shown that continuous target tracking in two and three dimensions can measure the time course of visual processing. We have also shown that millisecond-scale differences in visual processing between the eyes can be estimated with precision that is comparable to traditional psychophysics. Here, we discuss some of the advantages of the target tracking paradigm over traditional psychophysics, consider future methodological challenges, speculate about the potential for clinical application, and reflect on how the current results fit in the historical development of the science.

### Time-course information provides potential for enhanced explanatory power

The ability to recover the time course of information processing provides the potential for new explanatory power. For example, in the forced-choice psychophysics task, when sinusoidal target motion was presented in the plane of the screen and one eye was darker than the other, rather than the expected trajectory (Fig. 10A, black), one observer spontaneously reported perceiving elliptical trajectories in depth that were not aligned with the screen (Fig. 10A, colors). Upon follow-up debriefing, two additional observers reported perceiving misaligned elliptical trajectories.

**Figure 10.**
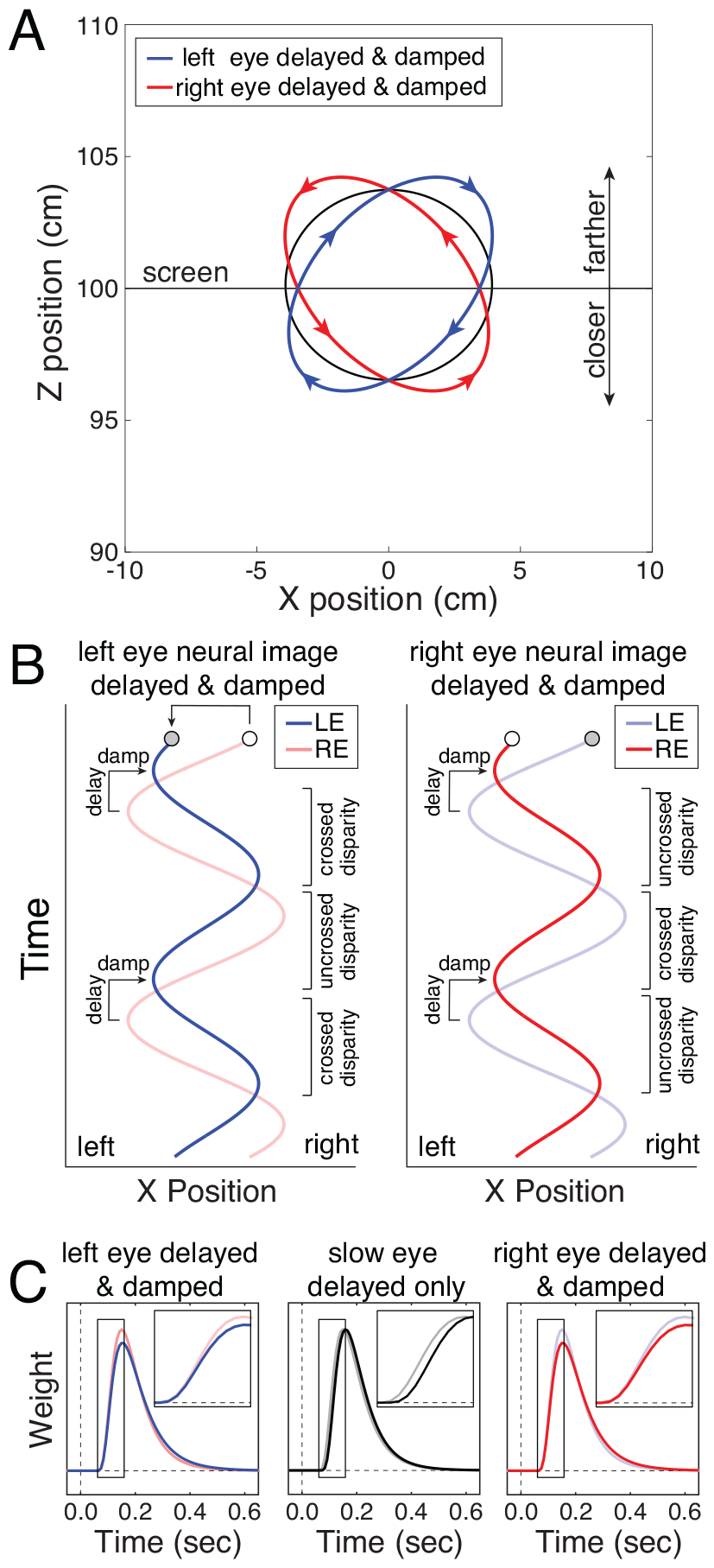
Anomalous Pulfrich percepts. **A** Some observers spontaneously reported perceiving anomalous near-elliptical motion trajectories that were not aligned with the screen (blue or red). Other observers perceived trajectories that were aligned with the screen (black). **B** Effective neural image positions as a function of time that can elicit the anomalous perceived motion trajectories in A. Left-eye (blue) and right-eye (red) neural image positions when the processing is delayed and damped in the left eye (left subplot) or right eye (right subplot). (Note that color schemes are consistent across all subpanels.) **C** Temporal impulse response functions that have different temporal integration periods in the two eyes can account for neural image positions in B. Impulse response functions with longer temporal integration periods tend to dampen the amplitude of the effective image position in that eye. Left subplot: Right-eye processing is fast and left-eye processing is delayed (i.e. has a longer time to peak) with a longer temporal integration period (i.e. has a larger full width at half-height). Middle subplot: Slow-eye processing is delayed but not damped relative to the fast-eye processing, and so will not cause anomalous Pulfrich percepts. Right subplot: Left-eye processing is fast and right-eye processing is delayed with a longer temporal integration period.

The most likely proximate cause of these anomalous Pulfrich percepts is that the amplitude of the effective image motion in the more slowly processed eye is not merely delayed but also damped relative to the effective image motion in the other eye (Fig. 10B). Shape differences between the impulse response functions in the two eyes can lead to such interocular differences in the amplitude of the effective neural image positions across time (Fig. 10C). Consider the case where the left eye is darker than the right eye, and the left-eye impulse response function peaks at a longer latency and has a longer temporal integration period than the right-eye impulse response function (Fig. 10C, left). The effective position of the left-eye image will be delayed and damped relative to the right-eye image (Fig. 10B, left). Stereo-geometry dictates that, as a consequence, observers will perceive the target stimulus undergoing ‘front left’ motion along a trajectory that is rotated such that its primary axis is askew to the screen (Fig. 10A, blue).

These ideas have recently been tested empirically. However, rather than using luminance differences between the eyes to induce temporal differences in visual processing, spatial frequency differences were used (Chin & Burge, 2022). High spatial frequencies reliably cause i) more delayed neural and psychophysical responses than low spatial frequencies (Bair & Movshon, 2004; Harweth & Levi, 1978) and ii) temporal integration periods with longer durations than low frequencies (Bair & Movshon, 2004; Vassilev, Mihaylova, & Bonnet, 2002). Traditional forced-choice and continuous target-tracking psychophysical experiments were conducted.

In the forced-choice experiment, observers viewed dichoptically-presented oscillating Gabors. The left- and right-eye Gabors were identical in all respects except that the carrier spatial frequency was higher in one eye than in the other (e.g. 3cpd in one eye, 1cpd in the other). The task was to report whether the apparent trajectory in depth was oriented left-side or right-side back from the plane of the screen (Fig. 10A; red and blue). The prediction was that the eye with the higher spatial frequency would have a longer temporal integration period (Fig. 10C). This longer temporal integration period would cause increased neural damping (Fig. 10B). And the increased neural damping would, in turn, cause the apparent motion trajectory to be misaligned with the plane of the screen (Fig. 10A). This prediction was borne out in the data. However, the evidence that the perceptual effects were due to temporal integration periods was quite indirect.

In the continuous psychophysics experiment, participants tracked Gabor targets that were identical to those used in the forced-choice experiment. The prediction was that the higher frequency Gabors would be associated with cross-correlograms having longer temporal integration periods (Fig. 10C). This prediction was also borne out in the data. And it was shown that the durations of the temporal integration periods estimated directly from the tracking experiment correlated well with the durations that were inferred from the forced-choice experiment. Continuous psychophysics thereby provided more direct evidence than traditional psychophysics for the proposed explanatory account of the anomalous Pulfrich effect (Fig. 10).

Hence, the results from the current article and from Chin & Burge (2022) together provide evidence that traditional and continuous psychophysics paradigms yield similar estimates of i) processing delay (or latency), and ii) the duration of temporal integration. The rich temporal information provided by continuous psychophysics can, in some cases, provide more direct evidence of the underlying causes of perceptual phenomena than can traditional psychophysics. Traditional psychophysics typically provides snapshots of visual processing and perception, whereas continuous psychophysics provides something more akin to video clips.

### Preservation of visual processing differences in the visuo-motor response

The millisecond-scale differences in processing that underlie the Pulfrich effect have almost certainly arisen by early visual cortex (Carney et al., 1989; Vassilev, Mihaylova, & Bonnet, 2002; Wolpert, Miall, Cumming, & Boniface, 1993). Indeed, there is a widespread view that these processing differences underlying the Pulfrich effect have their physiological origin in the retina itself (Mansfield & Daugman, 1978; Prestrude, 1971; Wolpert et al., 1993); also see (Bernhard, 1940)). We have shown that temporal delays in visual processing are faithfully preserved in the movement dynamics of the hand. To cause a motor response, electrical impulses must travel down multiple myelinated axon sheaths, and chemical communication must occur at multiple synaptic junctions as signals move from cortex, to the brain stem, and down the arm to the hand; the motion of the hand represents the culmination of all of these processes. Despite the myriad signal transformations and delays that occur after early visual cortex, the tiny interocular sensory delays that cause the Pulfrich effect are preserved in the motor response.

The fact that these small differences are preserved in the visuomotor response of the hand is striking. There is a clear evolutionary advantage for motor responses to occur as rapidly and consistently as visual processing allows. But there is no guarantee that this will happen. For example, the delay associated with a signal at one stage of processing may be magnified as the signal proceeds through subsequent stages of processing (Sternberg & Knoll, 1973). The fact that measurements from continuous target-tracking psychophysics is sensitive to small changes in temporal processing bodes well for its applicability to a wide array of research questions that depend on precise temporal information.

### Implications and applications

Traditional forced-choice psychophysical techniques are the gold standard in sensory-perceptual experiments. Forced-choice paradigms are our current best tools for minimizing measurement error. and they also sit atop a strong theoretical foundation for data analysis that is grounded in signal detection theory (D. M. Green & Swets, 1966). The price one pays for using traditional psychophysics, however, is that experimental data take a long time to collect, and the experiments themselves can be tedious for observers to participate in. While merely inconvenient in many laboratory settings, time and tedium become very real problems if one wants to make measurements in children or in clinical populations that might be unwilling or unable to produce a satisfactory number of trials (Candy & Cormack, 2022). Moreover, instructions that seem simple for experienced psychophysical observers, such as “press the left arrow key when the stimulus appears to be moving left when it is in front of the screen, and the right arrow if it appears to be going the other way” might be confusing to young children, and unintelligible to pre-verbal infants. Indeed, because infants naturally follow moving visual targets, continuous psychophysics should enable the behavioral assessment of visual function early in the developmental arc (Mestre, Neupane, Manh, Tarczy-Hornoch, & Candy, 2023). Finally, if one wishes to study effects that have large individual differences, then a technique is needed in which it is possible to obtain a satisfactory amount of data from a single observer in a short amount of time so that it is, in turn, possible to collect good data from a very large number of observers.

In clinical settings, vision is often assessed imprecisely, in large part because of severe time restrictions. Multiple visual functions–acuity, binocular function, and color vision, for example–are assessed with different tests requiring different instructions. There may be advantages to having patients perform only a single task—target tracking—and to have the target defined by different levels of spatial detail, binocular disparity, or color on different tracking runs (e.g. Chin & Burge, 2022). It has been recently shown, for example, that the contrast sensitivity function can be rapidly measured using continuous psychophysics (Mooney, Alam, Hill, & Prusky, 2020). Certain pathologies—psychophysical, physical, developmental, etc.—are likely to have sensory-motor processing delays as an indicating factor. For example, visual processing delays associated with amblyopia have recently been measured using continuous psychophysics (Gurman & Reynaud, 2024). Given its accuracy, efficiency, and relative ease of use, continuous psychophysics may have substantial utility as a tool for clinical assessment.

The arms and hands are large, heavy, and sluggish. The eyes are smaller, lighter, and more responsive. The eyes thus have the potential to increase the fidelity of tracking performance. When the eyes are tracking random walk motion, for example, the onset latency of smooth pursuit eye movements tends to be in the range of 80-120ms (Mulligan, Stevenson, & Cormack, 2013; Tavassoli & Ringach, 2009). Although smooth pursuit latencies depend on the properties of the target being tracked, these latencies are consistently less than the 150-200ms onset latencies of the motor response of the hand in the current tracking experiments (see Fig. 3). Delays in neural processing of sensory information have been shown to determine the delays of smooth pursuit eye movements (Lee, Joshua, Medina, & Lisberger, 2016). Finally, there tends to be considerably less inter-subject variability with smooth pursuit than with hand movements (Tavassoli & Ringach, 2009). By using eye movements, it should be possible to adapt continuous target tracking methods for use in young children and animal models. Young children and many animals reflexively follow moving targets with their eyes, potentially obviating the need for verbal instruction (in the case of children) or extensive training (in the case of animals). Indeed, continuous target tracking with eye movement monitoring has been used to measure how macaques and marmosets process optic flow (Knöll, Pillow, & Huk, 2018), and to measure both vergence and accommodative responses in infants as young as five weeks old (Downey, Pace, Seemiller, Candy, & Cormack, 2017). Using eye movements instead of, or in addition to, hand movements as the response effector in continuous target-tracking experiments may therefore have a number of advantages, and these probable advantages will only increase as eye tracking technology continues to improve.

Binocular eye tracking, in particular, might have interesting applications for assessing temporal processing differences between the eyes. First, binocular target tracking is potentially more efficient than monocular tracking; a single binocular run can yield information about interocular processing differences, whereas at least two distinct runs (one for each eye) must be performed with monocular tracking. Second, with binocular tracking, the interocular comparison is performed automatically by the binocular visual system before the motor system is engaged. With monocular target tracking, the comparison is performed only post-hoc by the experimenter after the motor response has been collected in two different conditions. Third, binocular tracking eliminates the possibility that state changes (e.g. alertness, motivation) between runs could corrupt estimates of interocular delay. Finally, as with the Pulfrich effect, small changes in one eye’s processing (e.g. due to a pathology) might produce detectable interocular differences even though the change in the monocular response per se would be impossible to detect without some previously-recorded baseline. Of course, for these benefits to be fully realized, the reasons underlying the mismatches between the binocular tracking predictions and data (see Fig. 9) must first be understood. Nevertheless, assessing the efficiency of monocular vs. binocular target tracking for such purposes could be a useful direction for future work.

The continuously changing stimulus and response also afford new possibilities for the application of analytical methods from systems neuroscience—or control theory (Bonnen et al., 2015; Burge, Ernst, & Banks, 2008; Straub & Rothkopf, 2022)—to psychophysical data. In research on human perception and behavior, it is common to compare human performance to that of a normative model of the task (Burge, 2020; Burge & Jaini, 2017; Ernst & Banks, 2002; Geisler, 2011; Kording & Wolpert, 2004). But normative models of psychophysical tasks are rarely constructed to predict continuous responses over time. In systems neuroscience research, however, popular models for neural systems identification are designed to recover computational-level descriptions of how stimuli drive neural response over time. Such models (e.g. the generalized linear model; GLM) are commonly applied to continuous-time stimuli and responses. But they have rarely been applied to human psychophysical data, perhaps because human datasets collected with traditional forced-choice methods have insufficient time-course information for the models’ full statistical power to be realized (Knoblauch & Maloney, 2008; Macke & Wichmann, 2010; Murray, 2012). The continuous target-tracking paradigm, paired with recent developments linking normative models to human performance (Burge & Geisler, 2015; Chin & Burge, 2020; Kim & Burge, 2018; 2020), and to methods for neural systems identification (Burg et al., 2021; Burge & Jaini, 2017; Iyer & Burge, 2019; Jaini & Burge, 2017; Park, Archer, Priebe, & Pillow, 2013), provides an exciting direction for future research.

### The reverse Pulfrich effect

It has long been known that interocular differences in luminance cause interocular differences in processing speed; the darker image is processed more slowly (Lit, 1949; Pulfrich, 1922). More recently, it was discovered that interocular differences in blur also cause interocular differences in processing speed: the blurrier image is processed more quickly (Burge et al., 2019; Rodriguez-Lopez et al., 2020). Given that blur decreases contrast and given that decreases in contrast tend to decrease processing speed (Albrecht, 1995; Bair & Movshon, 2004; Levi, Harwerth, & Manny, 1979; Nachmias, 1967; Reynaud & Hess, 2017; Shapley & Victor, 1978; Vassilev et al., 2002), it may at first seem surprising that blur increases rather than decreases processing speed. However, it is also known that, compared to lower spatial frequencies, higher spatial frequencies are processed with longer delays (Albrecht, 1995; Bair & Movshon, 2004; Levi et al., 1979; Min, Reynaud, & Hess, 2020; Nachmias, 1967; Reynaud & Hess, 2017; Shapley & Victor, 1978; Vassilev et al., 2002), and temporal integration periods (Bair & Movshon, 2004; Chin & Burge, 2022). Hence, because blur reduces the contrast of high spatial frequencies (fine detail) more than low spatial frequencies (coarse detail) (Burge & Geisler, 2011; 2012; Campbell & Green, 1965; Navarro, Artal, & Williams, 1993), a blurrier image, having fewer fine details, is processed more quickly. Experiments have confirmed that this explanation accounts for the reverse Pulfrich effect (Burge et al., 2019; Chin & Burge, 2022).

Many stimulus properties impact the temporal aspects of visual processing: luminance, contrast, spatial-frequency, blur, color. The visual field location at which a given stimulus property is processed also impacts processing delay and the duration of temporal integration (Dyer & Burge, 2023; Kelly, 1984). The computational rules relating image properties to processing speed remain to be discovered. The tracking paradigm, because of the rich temporal information it provides, may be well suited for investigating this problem as well.

## Conclusion

In this paper, we used continuous target-tracking psychophysics to demonstrate that small (i.e. less than 10ms) delays in visual information processing, likely arising in the retina, are preserved throughout the entire sensory-motor loop. We showed that differences in visual processing can be recovered from the motor responses of the hand. Continuous psychophysics is well-positioned to become an indispensable tool in the experimental toolkit for measuring the temporal dynamics of visual processing, an important but understudied topic in vision research.

## Acknowledgements

This work was supported by NIH grant R01-EY028571 from the National Eye Institute and the Office of Behavioral and Social Science Research to JB, and startup funds from the University of Pennsylvania to JB. The authors thank Stephanie Shields for extensive comments on a draft version of the manuscript. The authors also thank Kathryn Bonnen for useful discussion.

## Author Contributions

JB conceived the project, collected and analyzed data, performed simulations, and wrote the paper. JB and LKC developed concepts and edited the paper.

## Supplement

**Figure S1.**
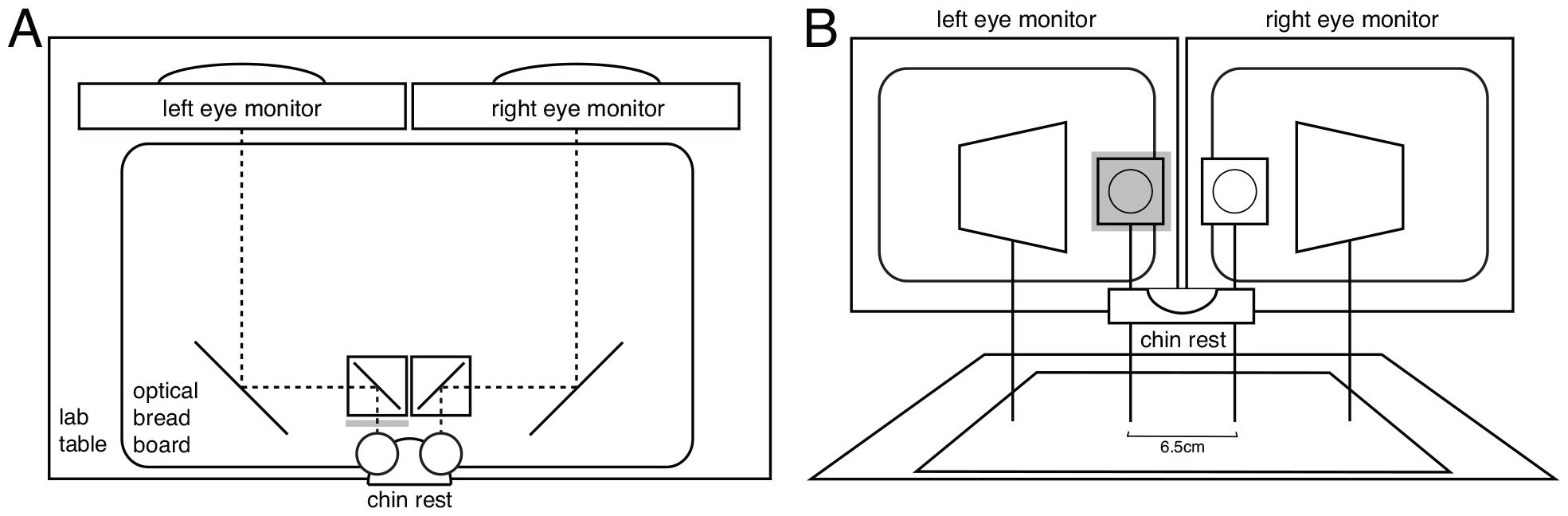
Haploscope apparatus for stimulus delivery. Each eye has its stimulus delivered by a dedicated monitor. The light bounces off two front-surface mirrors on its way to each eye (dashed lines). **A** Top down view. The dashed lines indicate the light path from the monitors to the eyes. A neural density filter is shown in front of the left eye. **B** Head on view. A neutral density filter is depicted in front of the left-eye viewport.

**Figure S2.**
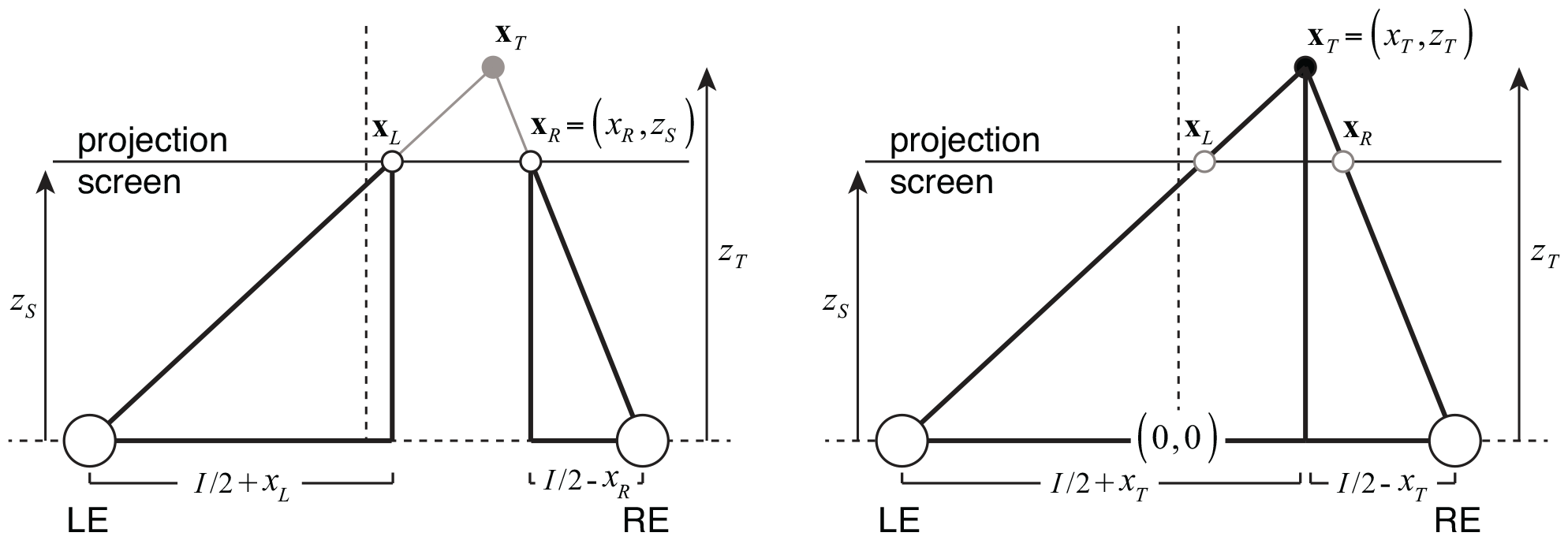
Projection geometry, similar triangles, and rendering positions. For a given target position x_*T*_ = (*x*_*T*_,*z*_*T*_) in stereoscopic space, the left- and right-eye stimuli must be rendered on the projection screen at positions **x**_*L*_ = (*x*_*L*_, *z*_*S*_) and **x**_*R*_ = (*x*_*R*_, *z*_*S*_), respectively. These positions are easily obtained using similar triangles (bold line segments in left and right subplots). Left-eye image position is obtained by solving the expression (*I / 2* + *x*_*L*_) (*I / 2* + *x*_*T*_) = *z* / *z*_*T*_ for *x*_*L*_ (Eq. 4a) where *I* is the interocular distance. Right-eye image position is obtained by solving the expression (*I* / 2 − *x*_*R*_) (*I* / 2 − *x*_*T*_) = *z*_*S*_ / *z*_*T*_ for *x*_*R*_ (Eq. 4b). In the actual apparatus, the left-eye stimulus and the right eye stimulus were displayed on separate monitors; but the geometry remains the same. Note that the drawings are not to scale.

### Supplementary Derivation

In the main text, we asserted in Eq. 1 that the difference between the left- and right-eye impulse response functions is proportional to the cross-correlation between the target velocity in X and the response velocity in Z

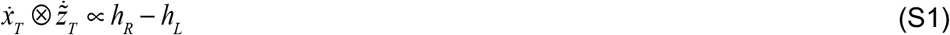

where 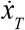 is the X velocity of the target, 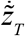 is the Z velocity of the human response, and *h*_*L*_ and *h*_*R*_ are the left- and right-eye impulse response functions, respectively. Here, we provide the derivation. Throughout the derivation (and in the main text), variables with dots denote velocities and variables with tildes indicate that they have been acted on by impulse response functions.

First, we substitute an expression for 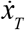 that has the form of Eq. 7a and substitute the expression for 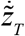 in Eq. 7b to obtain

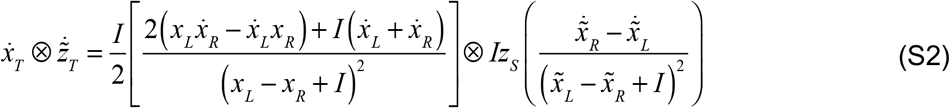

Given that the display screen was far from the observer (*z*_*S*_ =1000mm) and that the virtual position of the target in depth was never far from the display screen, the on-screen disparities (see Eq. 11) were always small relative to the interocular distance (i.e. a typical interocular distance *I* is 65mm, whereas 68% of the values assumed by *x*_*L*_ − *x*_*R*_ and 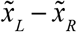 are between +1.2mm and 99% are between +3.9mm). Hence, (*x*_*L*_ − *x*_*R*_ + *I*) and 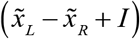 are approximately equal to *I*, allowing us to write

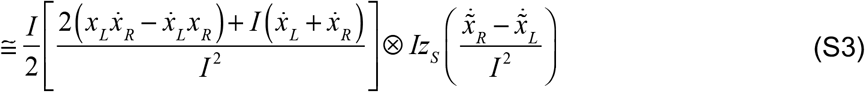

Grouping all the constant factors on the left-hand side and simplifying yields

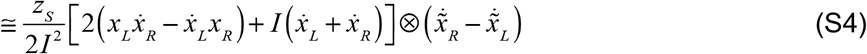

Now, note that the first parenthetical term in the square brackets has a positive and a negative term that are approximately equal because *x*_*L*_ and *x*_*R*_ are approximately equal to *x*_*T*_ and because 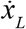 and 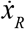 are approximately equal to 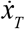. (Note that these approximate equalities again hold because the on-screen disparities were small.) These terms cancel, allowing us to write

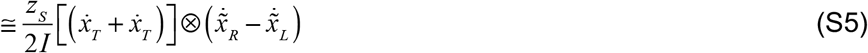

Expanding the second term by plugging in the effective left- and right-eye image velocities from the velocity equivalents of Eqs. 5a and 5b and simplifying gives

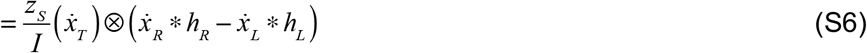

Substituting 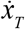 for 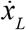 and 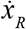, again making use of approximate equalities, gives

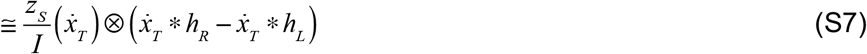

(Note that the above substitution ignores the fact that the rate of change of target disparity contributes to the Z response. But because the rate of the target X position is uncorrelated with the rate of change of disparity, it has no effect on the cross-correlation.)

Using the distributive property of convolution

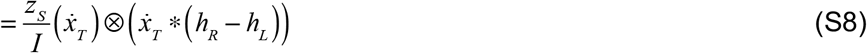

Using the associative property of cross-correlation and convolution

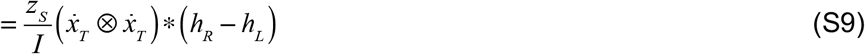

The autocorrelation of 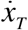 is a delta function centered at zero because the horizontal target velocities are distributed as white Gaussian noise by experimental design

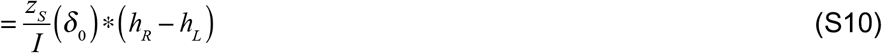

Convolving a delta function centered at zero with an arbitrary function yields the arbitrary function

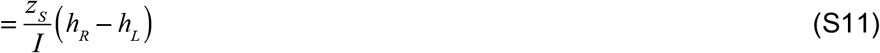

Finally, dropping the scale factors yields the proportionality asserted in the main text

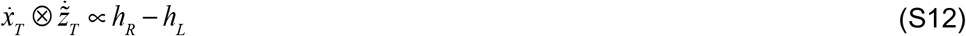

## Footnotes

1 Visuomotor tracking itself is not new. What is new is that visuomotor continuous target-tracking-psychophysics-based visuomotor measures of performance can be directly connected to measures of performance obtained from traditional forced-choice psychophysical techniques.

2 Visual and motor noise can reduce peak and average correlation values but should not impact the cross-correlogram shape (see Methods). Thus, noise should not bias estimates of delay or of the temporal integration period.

## References

Albrecht, D. G. (1995). Visual cortex neurons in monkey and cat: effect of contrast on the spatial and temporal phase transfer functions. Visual Neuroscience, 12(6), 1191–1210.

Bair, W., & Movshon, J. A. (2004). Adaptive temporal integration of motion in direction-selective neurons in macaque visual cortex. Journal of Neuroscience, 24(33), 7305–7323. 10.1523/JNEUROSCI.0554-04.2004

Banks, M. S., Gepshtein, S., & Landy, M. S. (2004). Why is spatial stereoresolution so low? Journal of Neuroscience, 24(9), 2077–2089. 10.1523/JNEUROSCI.3852-02.2004

Bernhard, C. G. (1940). Contributions to the neurophysiology of the optic pathway. Acta Physiologica Scandinavica, 1.

Bonnen, K., Burge, J., Yates, J., Pillow, J., & Cormack, L. K. (2015). Continuous psychophysics: Target-tracking to measure visual sensitivity. Journal of Vision, 15(3), 1–16. 10.1167/15.3.14

Bonnen, K., Huk AC, Cormack LK (2017). Dynamic mechanisms of visually guided 3D motion tracking. Journal of Neurophysiology, 118: 1515–1531, doi: 10.1152/jn.00831.2016.

Brainard, D. H. (1997). The Psychophysics Toolbox. Spatial Vision, 10(4), 433–436.

Burg, M. F., Cadena, S. A., Denfield, G. H., Walker, E. Y., Tolias, A. S., Bethge, M., & Ecker, A. S. (2021). Learning divisive normalization in primary visual cortex. PLoS Computational Biology, 17(6), e1009028. 10.1371/journal.pcbi.1009028

Burge, J. (2020). Image-computable ideal observers for tasks with natural stimuli. Annual Review of Vision Science, 6, 491–517. 10.1146/annurev-vision-030320-041134

Burge, J., Dyer, C. (2023). Eccentricity strongly modulates visual processing delays. bioRxiv, 559991, 1–18. doi: 10.1101/2023.09.30.559991

Burge, J., & Geisler, W. S. (2011). Optimal defocus estimation in individual natural images. Proceedings of the National Academy of Sciences, 108(40), 16849–16854. 10.1073/pnas.1108491108

Burge, J., & Geisler, W. S. (2012). Optimal defocus estimates from individual images for autofocusing a digital camera (Vol. 8299, p. 82990E). Presented at the Proceedings of the IS&T/SPIE 47th Annual Meeting, Burlingame, CA: Proceedings of SPIE. 10.1117/12.912066

Burge, J., & Geisler, W. S. (2014). Optimal disparity estimation in natural stereo images. Journal of Vision, 14(2), 1–18. 10.1167/14.2.1

Burge, J., & Geisler, W. S. (2015). Optimal speed estimation in natural image movies predicts human performance. Nature Communications, 6, 7900. 10.1038/ncomms8900

Burge, J., & Jaini, P. (2017). Accuracy Maximization Analysis for Sensory-Perceptual Tasks: Computational Improvements, Filter Robustness, and Coding Advantages for Scaled Additive Noise. PLoS Computational Biology, 13(2), e1005281. 10.1371/journal.pcbi.1005281

Burge, J., Ernst, M. O., & Banks, M. S. (2008). The statistical determinants of adaptation rate in human reaching. Journal of Vision, 8(4), 20.1–19. 10.1167/8.4.20

Burge, J., Rodriguez-Lopez, V., & Dorronsoro, C. (2019). Monovision and the Misperception of Motion. Current Biology, 29(15), 2586–2592.e4. 10.1016/j.cub.2019.06.070

Campbell, F. W., & Green, D. G. (1965). Optical and retinal factors affecting visual resolution. The Journal of Physiology, 181(3), 576–593.

Candy, T. R., & Cormack, L. K. (2022). Recent understanding of binocular vision in the natural environment with clinical implications. Progress in Retinal and Eye Research, 88, 101014. 10.1016/j.preteyeres.2021.101014

Carney, T., Paradiso, M. A., & Freeman, R. D. (1989). A physiological correlate of the Pulfrich effect in cortical neurons of the cat. Vision Research, 29(2), 155–165.

Chin, B. M., & Burge, J. (2020). Predicting the partition of behavioral variability in speed perception with naturalistic stimuli. Journal of Neuroscience, 40(4), 864–879. 10.1523/JNEUROSCI.1904-19.2019

Chin, B. M., & Burge, J. (2022). Perceptual consequences of interocular differences in the duration of temporal integration. Journal of Vision, 22(12), 12. 10.1167/jov.22.12.12

Cormack, L. K., Czuba, T. B., Knöll, J., & Huk, A. C. (2017). Binocular Mechanisms of 3D Motion Processing. Annual Review of Vision Science, 3, 297–318. 10.1146/annurev-vision-102016-061259

Cormack, L. K., Stevenson, S. B., & Schor, C. M. (1991). Interocular correlation, luminance contrast and cyclopean processing. Vision Research, 31(12), 2195–2207.

Cumming, B. G., & DeAngelis, G. C. (2001). The physiology of stereopsis. Annual Review of Neuroscience, 24, 203–238. 10.1146/annurev.neuro.24.1.203

DeAngelis, G. C., Ohzawa, I., & Freeman, R. D. (1991). Depth is encoded in the visual cortex by a specialized receptive field structure. Nature, 352(6331), 156–159. 10.1038/352156a0

Downey, C., Pace, G., Seemiller, E., Candy, T. R., & Cormack, L. (2017). Dynamic Characteristics of 5 to 22 week-old Infants’ Accommodation and Vergence Tracking Responses. Journal of Vision, 17(10), 443–443. 10.1167/17.10.443

Ernst, M. O., & Banks, M. S. (2002). Humans integrate visual and haptic information in a statistically optimal fashion. Nature, 415(6870), 429–433. 10.1038/415429a

Geisler, W. S. (2011). Contributions of ideal observer theory to vision research. Vision Research, 51(7), 771–781. 10.1016/j.visres.2010.09.027

Goldreich, D., Krauzlis, R. J., & Lisberger, S. G. (1992). Effect of Changing Feedback Delay on Spontaneous Oscillations in Smooth Pursuit Eye Movements of Monkeys. Journal of Neurophysiology, 67, 625–638.

Green, D. M., & Swets, J. A. (1966). Signal detection theory and psychophysics (Vol. 1). New York: Wiley & Sons.

Gurman, D., & Reynaud, A. (2024). Measuring the Interocular Delay and its Link to Visual Acuity in Amblyopia. Investigative Ophthalmology & Visual Science, 65(1), 2–2.

Harris, C. M., & Wolpert, D. M. (1998). Signal-dependent noise determines motor planning. Nature, 394(6695), 780–784. 10.1038/29528

Iyer, A. V., & Burge, J. (2018). Depth variation and stereo processing tasks in natural scenes. Journal of Vision, 18(6), 1–22. 10.1167/18.6.4

Iyer, A. V., & Burge, J. (2019). The statistics of how natural images drive the responses of neurons. Journal of Vision, 19(13), 4. 10.1167/19.13.4

Jaini, P., & Burge, J. (2017). Linking normative models of natural tasks to descriptive models of neural response. Journal of Vision, 17(12), 1–26. 10.1167/17.12.16

Julesz, B. (1964). Binocular depth perception without familiarity cues. Science, 145(3630), 356–362.

Kim, S., & Burge, J. (2018). The lawful imprecision of human surface tilt estimation in natural scenes. eLife, 7. 10.7554/eLife.31448

Kim, S., & Burge, J. (2020). Natural scene statistics predict how humans pool information across space in surface tilt estimation. PLoS Computational Biology, 16(6), e1007947–26. 10.1371/journal.pcbi.1007947

Knoblauch, K., & Maloney, L. T. (2008). Estimating classification images with generalized linear and additive models. Journal of Vision, 8(16), 1–19. 10.1167/8.16.10

Knöll, J., Pillow, J. W., & Huk, A. C. (2018). Lawful tracking of visual motion in humans, macaques, and marmosets in a naturalistic, continuous, and untrained behavioral context. Proceedings of the National Academy of Sciences, 115, E10486–E10494. 10.1073/pnas.1807192115/-/DCSupplemental

Kording, K. P., & Wolpert, D. M. (2004). Bayesian integration in sensorimotor learning. Nature, 427(6971), 244–247.

Kremers, J., Aher, A.J., Parry, N.R.A., Patel, N.B., Frishman, L.J. (2022). Electroretinographic responses to luminance and cone-isolating white noise stimuli in macaques. Frontiers in Neuroscience, 16:925405. doi:10.3389/fnins.2022.925405.

Lages, M., Mamassian, P., & Graf, E. W. (2003). Spatial and temporal tuning of motion in depth. Vision Research, 43(27), 2861–2873. 10.1016/j.visres.2003.08.006

Lee, J., Joshua, M., Medina, J. F., & Lisberger, S. G. (2016). Signal, Noise, and Variation in Neural and Sensory-Motor Latency. Neuron, 90(1), 165–176. 10.1016/j.neuron.2016.02.012

Levi, D. M., Harwerth, R. S., & Manny, R. E. (1979). Suprathreshold spatial frequency detection and binocular interaction in strabismic and anisometropic amblyopia. Investigative Ophthalmology & Visual Science, 18(7), 714–725.

Lit, A. (1949). The magnitude of the Pulfrich stereophenomenon as a function of binocular differences of intensity at various levels of illumination. The American Journal of Psychology, 62(2), 159–181.

Macke, J. H., & Wichmann, F. A. (2010). Estimating predictive stimulus features from psychophysical data: The decision image technique applied to human faces. Journal of Vision, 10(5), 1–24. 10.1167/10.5.22

Mansfield, R. J., & Daugman, J. G. (1978). Retinal mechanisms of visual latency. Vision Research, 18(9), 1247–1260. 10.1016/0042-6989(78)90111-6

Mestre, C., Neupane, S., Manh, V., Tarczy-Hornoch, K., & Candy, T. R. (2023). Vergence and accommodation responses in the control of intermittent exotropia. Ophthalmic & Physiological Optics, 43(4), 598–614. 10.1111/opo.13093

Min, S. H., Reynaud, A., & Hess, R. F. (2020). Interocular Differences in Spatial Frequency Influence the Pulfrich Effect. Vision, 4(20), 1–13. 10.3390/vision4010020

Mooney, S. W. J., Alam, N. M., Hill, N. J., & Prusky, G. T. (2020). Gradiate: A radial sweep approach to measuring detailed contrast sensitivity functions from eye movements. Journal of Vision, 20(13), 17. 10.1167/JOV.20.13.17

Morgan, M. J., & Thompson, P. (1975). Apparent motion and the Pulfrich effect. Perception-London-, 4(1), 3–18. 10.1068/p040003

Mulligan, J. B., Stevenson, S. B., & Cormack, L. K. (2013). Reflexive and voluntary control of smooth eye movements. In B. E. Rogowitz, T. N. Pappas, & H. de Ridder (Eds.), (Vol. 8651, pp. 86510Z1–22). Presented at the IS&T/SPIE Electronic Imaging XVIII, SPIE. 10.1117/12.2010333

Murray, R. F. (2012). Classification images and bubbles images in the generalized linear model. Journal of Vision, 12(7), 1–8. 10.1167/12.7.2

Nachmias, J. (1967). Effect of Exposure Duration on Visual Contrast Sensitivity with Square-Wave Gratings. Journal of the Optical Society of America, 57(3), 421–427.

Navarro, R., Artal, P., & Williams, D. R. (1993). Modulation transfer of the human eye as a function of retinal eccentricity. Journal of the Optical Society of America. a, Optics and Image Science, 10(2), 201–212.

Ogle, K. N. (1952). On the limits of stereoscopic vision. Journal of Experimental Psychology, 44(4), 253–259. 10.1037/h0057643

Ohzawa, I., DeAngelis, G. C., & Freeman, R. D. (1990). Stereoscopic depth discrimination in the visual cortex: neurons ideally suited as disparity detectors. Science, 249(4972), 1037–1041.

Park, I. M., Archer, E. W., Priebe, N., & Pillow, J. (2013). Spectral methods for neural characterization using generalized quadratic models. Advances in Neural Information Processing Systems, 1–9. Retrieved from http://papers.nips.cc/paper/4993-spectral-methods-for-neural-characterization-using-generalized-quadratic-models.pdf

Prestrude, A. M. (1971). Visual latencies at photopic levels of retinal illuminance. Vision Research, 11(4), 351–. 10.1016/0042-6989(71)90246-X

Pulfrich, C. (1922). Die Stereoskopie im Dienste der isochromen und heterochromen Photometrie. Die Naturwissenschaften, 10(35), 553–564.

Reynaud, A., & Hess, R. F. (2017). Interocular contrast difference drives illusory 3D percept. Scientific Reports, 7(1), 5587. 10.1038/s41598-017-06151-w

Rodriguez-Lopez, V., Dorronsoro, C., & Burge, J. (2020). Contact lenses can cause the reverse Pulfrich effect and anti-Pulfrich monovision corrections can eliminate it. Scientific Reports, 10:16086, doi: 10.1038/s41598-020-71395-y

Rodriguez-Lopez, V., Chin, B.M., & Burge, J. (2023). Decreases in overall light-level increase the severity of the reverse Pulfrich effect. bioRxiv, 559782, 1–17. doi: 10.1101/2023.09.27.559782

Rogers, B. J., & Anstis, S. M. (1972). Intensity versus adaptation and the Pulfrich stereo-phenomenon. Vision Research, 12(5), 909–928.

Shapley, R. M., & Victor, J. D. (1978). The effect of contrast on the transfer properties of cat retinal ganglion cells. The Journal of Physiology, 285, 275–298.

Sternberg, S., & Knoll, R. L. (1973). The perception of temporal order: Fundamental issues and a general model. In S. Kornblum (Ed.), Attention and Performance IV (pp. 629–685). New York: Academic Press.

Straub, D., & Rothkopf, C. A. (2022). Putting perception into action with inverse optimal control for continuous psychophysics. eLife, 11. 10.7554/eLife.76635

Tavassoli, A., & Ringach, D. L. (2009). Dynamics of smooth pursuit maintenance. Journal of Neurophysiology, 102(1), 110–118. 10.1152/jn.91320.2008

Tyler, C. W., & Julesz, B. (1978). Binocular cross-correlation in time and space. Vision Research, 18(1), 101–105.

Vassilev, A., Mihaylova, M., & Bonnet, C. (2002). On the delay in processing high spatial frequency visual information: reaction time and VEP latency study of the effect of local intensity of stimulation. Vision Research, 42(7), 851–864.

Watson, A. (1986). Temporal sensitivity. In K. Boff, J. Thomas, & L. Kaufman (Eds.), (Vol. 1). Handbook of perception and human performance.

Wheatstone, C. (1838). On some remarkable, and hitherto unobserved, phenomena of binocular vision. Philosophical Transactions of the Royal Society of London, 128, 371–394.

Wilson, J. A., & Anstis, S. M. (1969). Visual delay as a function of luminance. The American Journal of Psychology, 82(3), 350–358.

Wolpert, D. M., Miall, R. C., Cumming, B., & Boniface, S. J. (1993). Retinal adaptation of visual processing time delays. Vision Research, 33(10), 1421–1430.S

